# Towards a Health-Associated Core Keystone (HACK) index for the human gut microbiome

**DOI:** 10.1101/2024.05.27.596018

**Authors:** Abhishek Goel, Omprakash Shete, Sourav Goswami, Amit Samal, Lavanya CB, Saurabh Kedia, Vineet Ahuja, Paul W O’Toole, Fergus Shanahan, Tarini Shankar Ghosh

**Affiliations:** Department of Computational Biology, Indraprastha Institute of Information Technology (IIIT-Delhi), Okhla Phase III, New Delhi, India; Department of Gastroenterology, All India Institute of Medical Sciences (AIIMS), New Delhi, India; Schools of Microbiology and Medicine, APC Microbiome Ireland, University College Cork, Ireland

## Abstract

Variations in the normal gut microbiome and the existence of context-dependent disease associations have confounded the identification of microbiome markers of health. A reliable indexing of taxa based on their association with host health and microbiome resilience could accelerate development of microbiome-based therapeutics including selection of live biotherapeutics and facilitate microbiome comparisons in diverse study populations.

Here we first investigated 196 taxa for their association with three hallmark properties of health and microbiome-resilience, namely prevalence/community-influence in non-diseased subjects, longitudinal stability and host health, using a discovery cohort of 39,926 publicly available adult (> 18 years) gut microbiomes from 127 studies spanning 42 countries and 28 disease conditions (including 9,434 longitudinal samples). We identified 18 species-level-taxa, referred to as ‘Health-Associated Core Keystones’ (HACKs), with association-strengths in the top 30 percentile for all three properties. We integrated these association-strengths into a single value, the HACK-index, that ranks taxa based on their estimated contribution to both microbiome stability and host health. We then demonstrate the reproducibility of these indices for the taxon associations with the three properties, using a validation cohort of 4,500 gut microbiomes (from 11 studies with various demographics and diseases). Specific consortia of high HACK-index taxa are also associated with positive response to Mediterranean Diet interventions and Immuno-Checkpoint-Inhibitor therapies. We finally identify the distinguishing microbial genomic functions associated with high HACK-indices/HACK-taxa that can be investigated using targeted mechanistic studies to identify microbiome effectors of improved health.

The availability of HACK-indices provides a rational basis for microbiome comparisons and will facilitate the selection and design of microbiome-based therapeutics.

## Introduction

An evolving area of microbiome research is the development of gut microbiome-based therapeutics. This class of therapeutics aims to restore or modulate the gut microbiome to ameliorate diseases and improve human health. Strategies include ‘probiotic’ supplements (often *Lactobacilli* and *Bifidobacteria*) and live biotherapeutics (such as resident ‘beneficial’ microbes, currently led by *Faecalibacterium* or *Akkermansia*); prebiotics (dietary ingredients that promote ‘beneficial’ members of the gut community); or synbiotics (combinations of both); and faecal microbiome transplantation^1–3^. Since the success of these interventions varies in different study populations^4,5^, a consensus on what constitutes a health-associated stable gut microbiome is highly desirable but difficult to achieve.^6^

The adult human microbiome varies widely in both health and disease, and the interpretation of different studies is confounded by multiple factors including geographic region, socio-economic status, age, and personal life-style^6–8^. Moreover, some microbiome components exhibit context-dependent associations with diseases (e.g. *Akkermansia*)^9^. This suggests that restrictive labelling of species with terms like ‘good’, ‘bad’, ‘beneficial or ‘detrimental’ is inappropriate. However, since meta-analyses have identified groups of gut microbial community members to be commonly depleted or enriched across multiple diseases^7,10–14^ an alternative approach may be to consider the microbiome as a spectrum where different microbiome members are variably linked in ranked-order with host health. Species higher up in this order might have higher potential as microbial therapeutics or as targets (or clinical end-points) for enrichment by microbiome-based therapeutics.

An ideal taxonomic candidate for supplementation/enrichment in microbiome-based therapeutics should have two additional properties (besides a common association with health). It should have a positive linkage with long-term gut microbial stability^15–17^ and it should be a reasonably conserved ‘interactive’ (both an influencer and a responder to others) and core member of the non-diseased gut microbiome^11,18,19^. We hypothesized that a global ranking index of gut microbial taxa associated with these properties would provide a priority list of taxa for formulation as well as measuring the efficacies of microbiome-based therapeutics promoting general health. It is also computable using recently available cross-sectional and longitudinal gut microbiome sequence data from diverse population cohorts spanning a wide-variety of diseases.

Here, we generated for the first time a ranking of 196 gut microbial taxa - the ‘HACK Index’ (Health-Associated Core-Keystone Index) – by scoring these taxa for their association with three quantifiable properties: prevalence/community-influence in non-diseased subjects, longitudinal-stability and negative association with disease. The index was computed based on a global meta-investigation involving a discovery cohort of 39,926 gut microbiomes from 127 publicly available study cohorts (including both cross-sectional and longitudinally sampled datasets spanning 42 countries and 28 different disease conditions. Using a validation dataset of 11 additional study-cohorts comprising 4,500 gut microbiomes, we demonstrate the reproducibility of these indices, where taxa with higher HACK indices have significantly higher influence in the community and positive associations with health and microbiome stability. Notably, we also observe that specific taxa with high HACK indices are associated with favourable response to diverse microbiome-associated therapeutic interventions like the Mediterranean Diet and Immuno-Checkpoint Inhibitor therapy in Cancer. We also identify functional/metabolic functionalities associated with HACK scores, that can be studied to investigate mechanisms.

## Results

### Global ranking of gut microbial taxa based on their prevalence and community influence in non-diseased adult gut microbiomes

A key characteristic for a taxon linked with microbiome stability/host health is that it should be a prevalent member and have high level of community interaction in gut microbiomes from non-diseased individuals. We refer to this property as Core Influence (‘Core’ as these members are more prevalent; ‘Influence’ as they are expected to have strong association with the composition of the other members of the gut microbial community computed as described below). We predict that supplementation of the microbiome with such taxa would have a higher likelihood of transitioning it towards a normal-like microbiome.

The collated repository of 39,996 gut microbiomes contained data from primarily three kinds of datasets: population level studies on apparently non-diseased (‘control’) subjects; longitudinal study cohorts collecting samples from subjects (both diseased and non-diseased) at multiple time-points and thirdly; matched disease-control cohorts focusing on different diseases (**See Supplementary Text 1; Supplementary Table 1; Fig 1A; Methods**). For this phase of the analysis, we used a repository of 18,642 gut microbiomes from apparently non-diseased (or control) adult subjects (age >= 18 years) collected from 72 studies from 34 nationalities (**Supplementary Text 1**). These included taxon abundance profiles derived from both shotgun sequencing (n=9,736)^20–22^, and 16S rRNA amplicon sequencing projects (n=8,906) (**Supplementary Text 1**; **Supplementary Table 2**). This large collation of gut microbiome profiles from control study populations encompassed diverse life-style patterns ranging from industrialized/urban populations of Europe, North America and East Asia to rural and hunter-gatherers from Africa, India and South America; from population cohorts with a mix of rural and urban life-styles from India, China, Columbia, Fiji, El-Salvador, Mongolian and Oceania to populations with transitional life-styles like the Irish Travellers.

**Figure 1.**
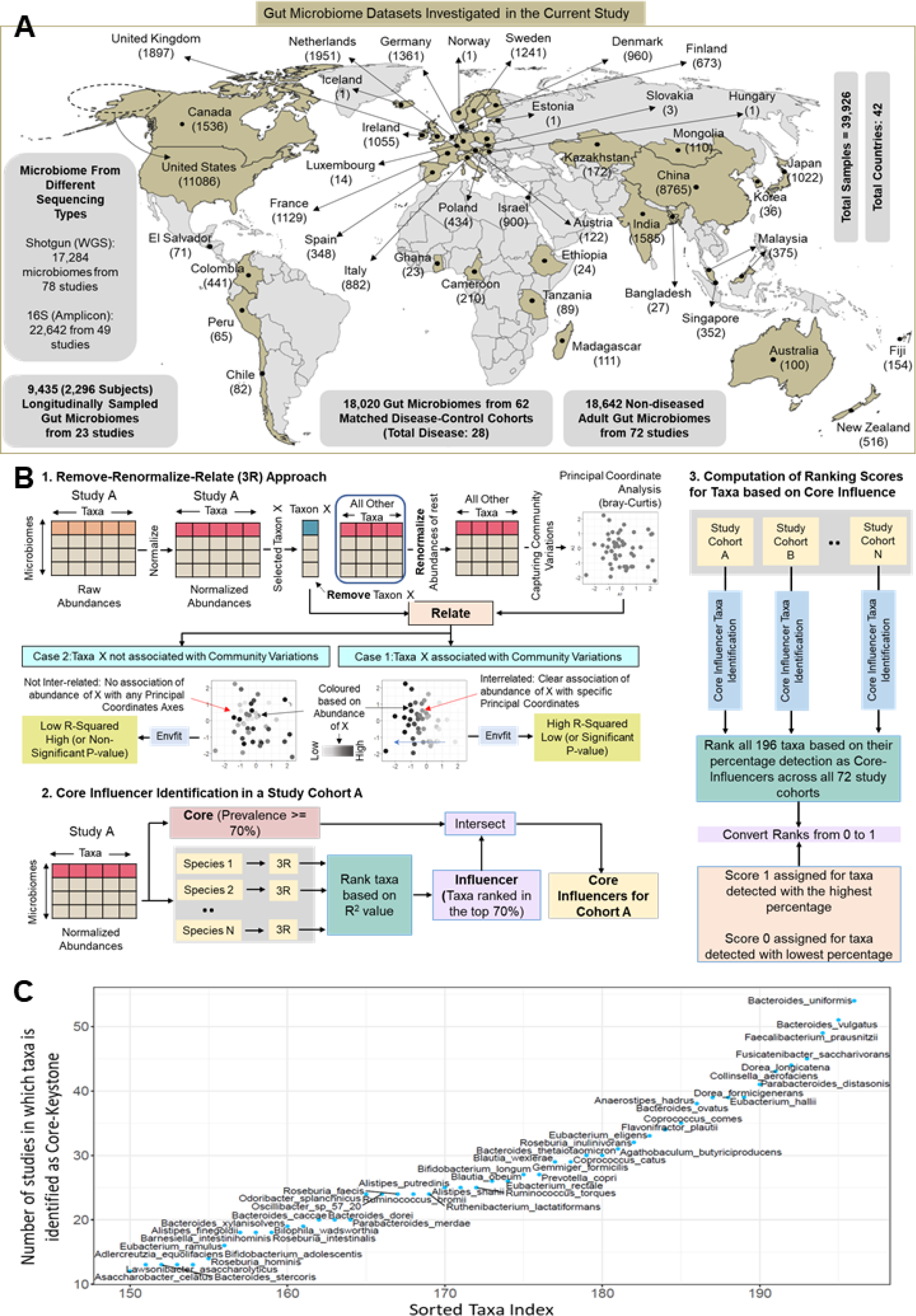
Global range of the microbiome data investigated in this study along with the computation of the ranked order of Core-Influences across the 72 study-cohorts comprising non-diseased adult subjects. A. Global country-wise distribution of the 39,926 microbiomes investigated in the current study. Also indicated are the number of samples or microbiomes belonging to each of the sequencing types (WGS and 16S) and the different number of microbiomes along with the number of studies considered for the three different types of investigations, performed in this study. B. Schematic description of the Core-Influencer taxa identification frame-work adopted for each study using the prevalence + Remove-Renormalize-Relate (3R) approach (described in top panel). Briefly, to measure the community influence of a given taxon (X) in a study cohort A, we first removed the abundances of X from all microbiomes of this cohort and recomputed the abundances by renormalizing the abundances of the remaining taxa. Community variations of the resulting microbiome profiles were then computed using Principal Coordinate Analysis (PCoA). Subsequently, these community variations were related with the abundance of X using the envfit approach. In scenarios where the abundance of X is strongly associated with the compositions of all other members in a gut microbial community, community-wide variations as measured using specific Principal Coordinates (or PCoA axes) will show a strong correlation (computed using the envfit approach) with the abundance of X (as shown in this representative example). The 3R analysis was applied to all 196 taxa across a given study and taxa were ranked based on their R-squared values. Taxa with R-squared in the top 70% were selected. In parallel, we also identified the prevalent taxa in the dataset (70% prevalence) (Bottom panel). Taxa that satisfied both the properties were then selected as the Core Influencers f the study cohort. C. Ranked order of gut microbial taxa arranged in decreasing order of number of study cohorts in which they were identified as a Core Influencer. This provides the ranking scores for the different taxa based on the consistency of their detection as Core Influencers across the 72 study cohorts.

For this and the subsequent investigations, we specifically selected a list of 196 species-level taxa that were detected in >= 10% samples in >= 35% of the study cohorts (‘Gut-Associated’). Cumulatively, these taxa accounted for a median abundance of 97.7% across all 18,642 gut microbiomes from 72 studies (**Supplementary** Figure 1A). Both properties of prevalence and influence are quantifiable. The prevalence of a given taxon X was calculated as the percentage of microbiomes from a study cohort in which the taxon is detected. For calculating the influence, we utilized a modified version of the recently developed Empirical Presence-absence Interrelation (EPI) approach (that enables quantifiable prediction of the degree of community influence of each microbiome member by measuring the community-wide differences in microbiomes in the presence/absence of the taxon)^23^. This modified version, referred to as the Remove-Renormalize-Relate (3R) approach, computed the community influence (or association) of a given taxon for a given study cohort by correlating its abundance variation (rather than its detection as in EPI) with the community-wide variations of the remaining taxa across all gut microbiomes of this cohort (**Fig 1B; Methods**). The approach was based on the premise that the stronger the associations of X with other members of a community, the more significantly will its abundance/representation in the community correlate with the community-wide variations of all other taxa.

Implementing the 3R approach had three stages: first, to ensure that the estimated variations in community composition were not simply due to large variations in the relative abundances of X, but were solely based on the differences in the mutual representation of remaining members, we removed X from the community abundance profiles and renormalized the abundances of the remaining members; next, we profiled the variations of the resultant community-taxonomic profiles of all gut microbiomes of the given cohort using Principal Coordinate Analysis (PCoA); finally, the envfit approach was utilized to correlate the abundance of X with the community-wide variations for the cohort (higher R^2^ of the envfit approach indicating higher concordance between abundance of X and the PCoA-profiled community-wide variations) (**See Methods; Fig 1B**). In other words, the higher the envfit-R^2^ the greater is the likelihood that microbiomes with similar representations/abundance of X will tend to have lower variations in their community composition or occupy similar positions in the PCoA space and *vice-versa* For each study, the above steps were repeated for all 196 taxa (**see above**). Taxa were then ranked based on their extent of community influence (envfit-R^2^) (using the 3R approach; **see Methods, Fig 1B**). Taxa with ranks >=70% and also detected in >= 70% samples of a given study cohort were identified as the Core-Influencer taxa for that cohort. This process was repeated for all study cohorts and taxa were ranked based on their consistency of detection as Core-Influencer taxa across the 72 study cohorts (**Supplementary Table 3)**. This produced a first ranked order of taxa with respect to the first probed hallmark of microbiome health, that of high prevalence and high community Influence in non-diseased subjects (**Supplementary** Figure 2; **Fig 1C**). For each taxon X, we thus calculated the Core- Influence score (CS) as the percentage of cohorts for which the taxon was identified as a ‘Core- Influencer’.

The majority (76%) of the species tested (149) did not show any strong Core-Influencer properties (i.e. not detected as Core-Influencer-Taxa in > 80% of studies). The remaining 47 taxa formed a gradient with increasing consistency of prediction across cohorts (**Fig 1C**). These 47 taxa comprised greater than 70% of the list of Core-Influencer taxa identified in greater than 80% of the study cohorts. This indicates that the Core-Influencer property is restricted to a small subset of the gut microbiome. The top 5 ranked taxa (in terms of their consistency of detection as Core-Influencer taxa) consisted of two *Bacteroides* species, followed by *Faecalibacterium prausnitzii*, *Fusicatenibacter saccharivorans* and *Dorea longicatena*. While many of these taxa have been associated with health in prior studies^11,24^, others have been associated with faster gut microbiome recovery after antibiotic treatment^25^ and resilient donor microbiome colonization following faecal microbiome transplantation^26^. Notably, the 47 top predicted Core Influencer taxa also included members like *Flavonifractor plautii* and *Collinsella aerofaciens*, previously shown as enriched across multiple diseases^7^. Thus, the Core-Influencer taxa while being associated with high community influence and prevalence in the microbiome of apparently “healthy” subjects, may not be directly linked to health. Furthermore, across the cohorts, the Core-Influencer property was not directly related to species abundance because simply being highly abundant did not necessarily qualify a taxon to be a Core-Influencer-Taxa of a normal gut microbiome (**Supplementary** Figure 3).

Furthermore, while the detection pattern of Core-Influencers was not significantly influenced by the sequencing-type of study cohort (16S or WGS) (PERMANOVA R-Squared: 0.01, P-value: 0.46) (**Supplementary** Figure 4), it was significantly influenced by the Cohort-Lifestyle patterns (specifically the extent of urbanization; profiled by categorizing the study-cohorts into three ‘CohortTypes’, namely namely ‘IndustrializedUrban’, ‘UrbanRuralMixed’ and ‘RuralTribal’, based on the predominant life-style patterns of the study populations) (PERMANOVA: P-value = 0.001) (**Supplementary** Figure 5) and commensurate with the variations in the overall microbiome compositions (Procuste Analysis: R=0.72; P=0.001) (**Supplementary Text 2**; **Supplementary** Figure 5). Reflecting observations from other studies, multiple Bacteroides species and Prevotella copri showed noticeably higher incidence as Core Influencers for the IndustrializedUrban and Rural/Tribal (and Mixed) cohort groups, respectively. We also observed previously unreported life-style specific assocations for other taxa **(Supplementary** Figure 6).

To further evaluate the confounding effect of these cohort-life-style specific variations on the overall Core Influence scores, we then computed CohortType-specific Core-Influence scores (three separate Core-Influence scores using the above 3R+Prevalence approach separately for study-cohorts of each CohortType) and then computing the coefficient of variation of these scores for each of 196 taxa across the Cohort-Types (**Supplementary Text 2**). Notably, this coefficient of variation of Core-Influence scores (across Cohort-Type) showed a significant negative correlation with overall Core-Influence scores calculated by considering all 72 cohorts (irrespective of CohortType) (Spearman Rho = -0.77, P=2.2e-16) (**Supplementary** Figure 7). This indicates that taxa with higher overall Core-Influencer scores show significantly lower variations in their Core-Influence properties across cohorts belonging to different life-styles.

### Meta-analytic investigation linking taxon abundance with intra-subject microbiome variability to rank taxa based on their association with temporal stability

The Core Influence indicates the efficiency of a taxon for colonizing the gut and interacting with the other gut microbial community members of non-diseased subjects. However, it does not indicate whether its removal has any enduring effect on the composition of the gut microbiome. From the perspective of microbiome modulation or restoration, it is important to not only identify taxa that are prevalent and interactive, but also those which are associated with stability. For this purpose, we performed a second investigation for the association of the different taxa to intra-subject variability, an inverse measure of longitudinal microbiome stability.

For this, we used 23 distinct study cohorts (**Supplementary Table 1**), encompassing 9,435 longitudinally-sampled gut microbiome samples, from 2,243 subjects. Of these datasets, 7,134 microbiome profiles contained at least one follow-up sample from the same subject at a subsequent time-point. The study cohorts included in this investigation contained data from different kinds of subjects (including non-diseased controls, patients with ulcerative colitis, Crohns’ disease and irritable bowel syndrome and subjects treated with different antibiotics and fibre interventions; See **Methods**). A stable microbiome is expected to display low temporal variability across consecutive time-points^22^. If a specific gut-associated taxon is associated with microbiome stability, higher abundance of this taxon in a microbial community sampled at time ‘t’ is likely to be associated with lower alterations in composition of the same community when sampled at a subsequent time-point ‘t+1’ and, vice-versa. To test this, for each of the 7,134 gut microbiomes with follow-ups, we first computed a ‘Follow-up Distance’ as the distance between the species-level microbiome composition of a given microbiome with that collected from the same subject at the next subsequent time-point (**Methods; Fig 2A**). The Follow-up distance thus is an inverse indicator of microbiome stability. We computed Follow-up distances using two different distance measures, Bray-Curtis and Aitchison. Next, we investigated the association of each taxon by correlating their abundances in each microbiome with the corresponding Follow-up distances using an iterative Random-Effects model based meta-analysis approach (**Fig. 2A and Methods for a detailed explanation**). The approach consisted of 10 iterations, where in each iteration, the 196 gut-associated taxa were provided a stability-association score based on the strength and consistency of their positive association with temporal microbiome variability across the study cohorts (separate scores computed for either distance measures and then averaged; described in **Fig 2A**; **Supplementary Table 4)**. The final stability-association score (SS) for each taxon was then computed as the global mean of all stability-association scores obtained for the taxa across iterations for both distance measures. The SS provided the second ranked order of taxa based on their links with another hallmark, longitudinal microbiome stability (**Supplementary Table 5**).

**Figure 2:**
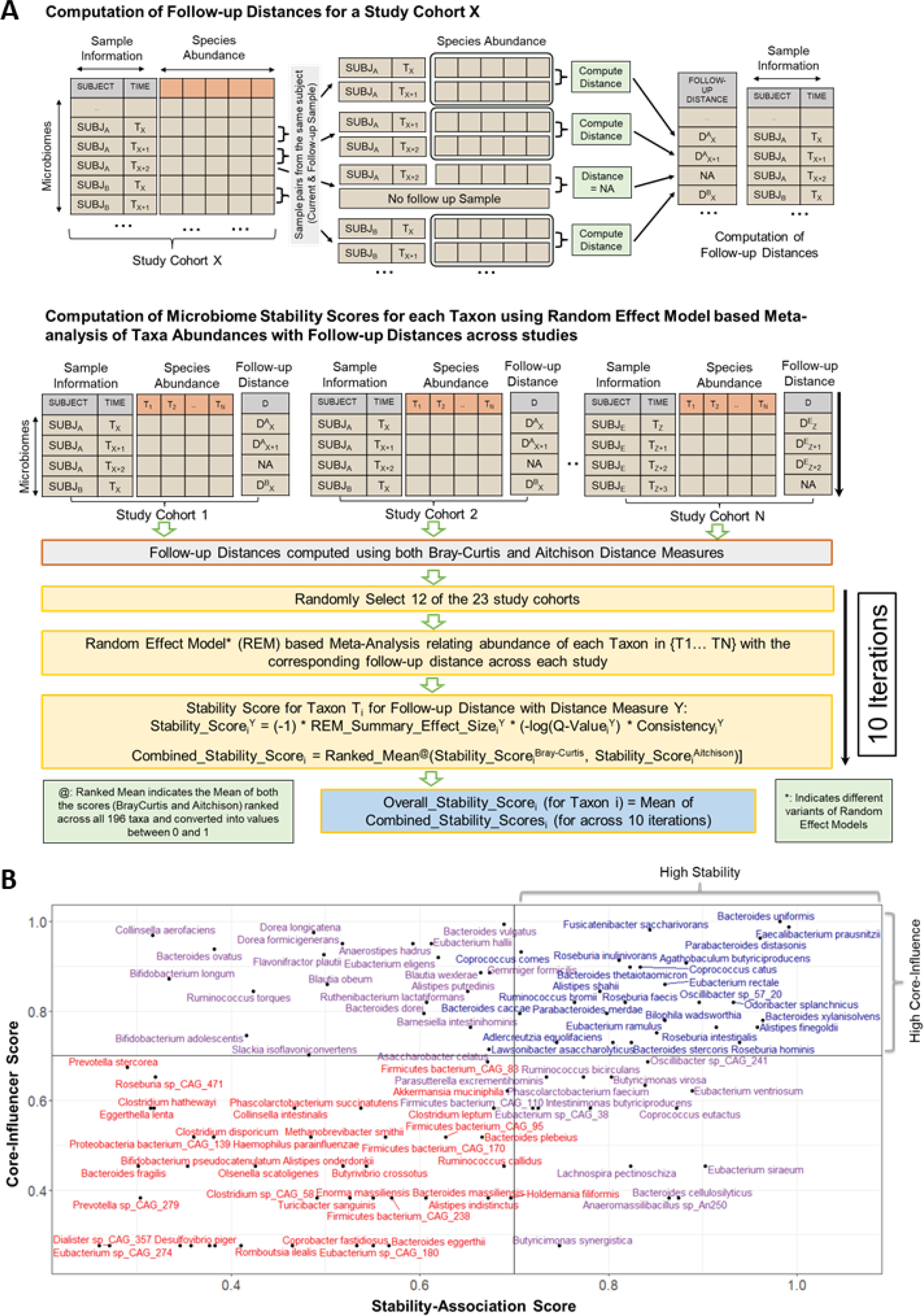
**Computation of Stability-Association Scores using a meta-analysis based investigation of 23 longitudinal study cohorts**. A. Schematic description of the methodology to compute the Stability-association score of each taxon. The method was based on the computation of Follow-up distances (described in the top panel). Follow-up distance was the distance between the microbiome from a given subject at a given time-point and that collected from the same subject at the immediately subsequent time-point using any distance measures (we used Aitchison and Bray-Curtis). To rank taxa for their association with microbiome stability, we performed 10 iterations. In each iteration, we selected a subset of 12 studies and performed Random Effects Model based analysis computing the association of each taxon with the two Follow-up distances. In each iteration, for a given distance measure, we computed stability-association scores for each taxon by multiplying the magnitude of the summarized estimate (multiplied by -1 as Follow-up distance is an inverse indicator of stability), the significance (-log10 of q-value) and the consistency (See Methods) of its abundance association with the Aitchison and Bray-Curtis Follow-up distances across cohort. Separate scores for the two distance-measures were then averaged to get a final iteration-specific stability-association score. This iterative boot-strapped based study-selection approach enabled us to reduce the influence of individual studies with possibly atypical effects. The final stability-association score (SS) was then calculated as the overall mean of the iteration-specific stability-association scores. **B.** Plot grouping the different taxa based on their Core-Influencer Scores (obtained in Fig 1) and the Stability-Association scores obtained. We identify a group of 26 taxa having both Core-Influencer score and Stability Score in the top 70%. We refer to these as the group of Core Keystones.

We identified 26 taxa having both ranking scores (CS and SS) in the top 70% (**Fig 2B**). These included *Faecalibacterium prausnitzii*, *Bacteroides uniformis*, *Bacteroides xylanisolvens*, *Odoribacter splanchnicus*, *Roseburia hominis*, *Roseburia inulinivorans*, *Coprococcus catus*, *Coprococcus comes*, *Agathobaculum butyriproducens*, *Fusicatenibacter saccharivorans* and *Eubacterium rectale*. We refer to this set of 26 taxa as the set of Core Keystones (Core: because they were prevalent; Keystone: because they had high influence and were associated with microbiome stability).

We also performed additional validation to ensure that the obtained stability-association scores were not majorly confounded by the variabilities of clinical phenotype of the participants of the different longitudinal cohorts, and were not noticeably affected by the across-time-point abundance variations of a taxon itself (**Supplementary Text 3**; **Supplementary** Figure 8**; Supplementary Tables 6-11**). Stability-association scores computed across each of the subject groups and abundance-adjusted variants with the overall Stability-Association scores computed in **Fig 2A** (ranging from 0.83 to 0.53) (**Supplementary** Figure 8). This suggests that the Stability-Association score robustly captured the overall rank-order of taxon associations with microbiome stability individually within the different subject groups.

### Ranking taxa based on their association with health

The final property to be examined was the association of taxa with health. For this investigation, we selected 18,020 microbiomes from 62 study cohorts (**Supplementary Table 12**). These were selected in a step-wise manner by ensuring that each cohort had sufficient samples from the diseased and non-diseased groups and was not heavily biased (in terms of sample numbers) towards any specific group. This selected set covered 66 different diseases/conditions that could be further categorized into 28 different disease categories (**Supplementary Text 1**; **Supplementary Tables 13-14**). The next step was the identification of markers of health and disease across the 28 disease-categories. All cohorts corresponding to a given disease category were first grouped (this was done to provide equal weighting to all disease categories without any bias towards disease groups having higher number of study cohorts). Subsequently, we identified the disease- and disease-negative taxa corresponding to each using Mann-Whitney tests (**Fig 3A**).

**Figure 3.**
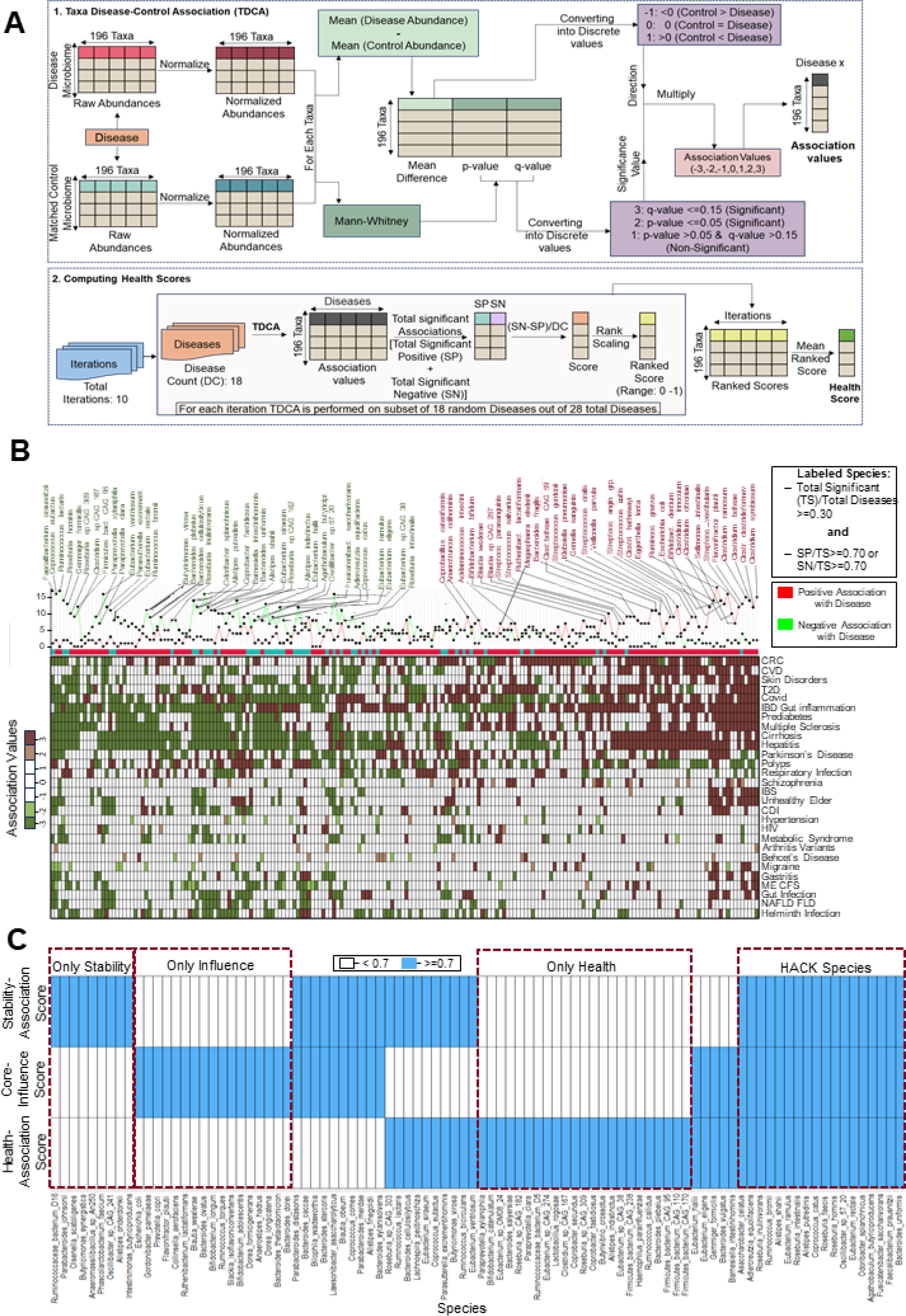
Computation of Health-Association Scores of the 196 taxa (based on the consistency of their alterations across 28 disease categories), and the identification of different taxa groups based on their association with the three different scores. A. The top panel describes the Taxon Disease-Control Association (TDCA) approach adopted for identifying the association (or alteration) patterns of the 196 taxa with the different Disease categories. Each taxon was assigned an association tag based on the directionality of the change and the significance of the change (p-value using Mann-Whitney tests and corrected Q-values computed using these p-values and the Benjamini-Hochberg FDR approach) (as summarized in **B**). The bottom panel describes the iteration-based approach used for computing the final Health-Association scores for the 196 taxa. We performed 10 iterations, each time selecting 18 of the 28 disease categories, and identified the directionality of taxa-associations with each of the selected disease categories. Each taxon was subsequently assigned a score, computed as the difference between the number of disease categories where it was significantly (Q<=0.15) depleted (SN) minus the number of categories it was enriched (SP) divided by the total number of disease categories (DC). This score was ranked across taxa to obtain the iteration-specific Health-Association scores. Scores were then averaged across the 10 iterations to yield the final Health-Association score for each taxon. **B.** The lower heatmap presents the association values (computed as in **A**) for the 196 taxa across the 28 disease categories. Columns indicate taxa and rows show the Disease categories. The two line-plots on the top panel shows the total count of disease categories where the corresponding taxon is significantly enriched (red-line) and depleted (green-line). Shown on top are taxa, which were consistently and significantly (Q- value <= 0.15) either enriched (shown in red font) or depleted (green font) across at least 30% of the considered disease categories. **C.** Heatmap showing the pattern of the 93 taxa showing association scores in the top 30% for at least one of the three properties. The grouping of different taxa based on their association patterns is also indicated.

There were considerable variations in the overall number of taxon alterations across the 28 different disease categories, with some diseases like Colorectal Cancer (CRC), cardiovascular diseases (CVD), Prediabetes, Type-II-Diabetes, Covid, IBD and other gut inflammations, especially pronounced with respect to the number of microbiome alterations (**Fig 3B**). Nevertheless, as previously noted in multiple meta-analyses^7,11,12,24^, certain taxa exhibited reasonable consistency in their alterations across different diseases (**Fig 3B**). Of these, 37 (health-associated) were consistently depleted (Q-value <= 0.15) and 33 (disease-associated) were consistently enriched (Q-value <= 0.15) (**Supplementary Table 15**). Amongst the health-associated taxa, those with strongest associations were *Ruminococcus lactaris*, *Faecalibacterium prausnitzii*, *Eubacterium ventriosum*, *Barnesiella intestinihominis*, *Eubacterium eligens*, *Coprococcus eutactus* (depleted in <= 50% disease categories). At the other end of the spectrum were the strongest disease-associated taxa, starting from *Ruminococcus gnavus*, *Clostridium symbiosum*, *Clostridium bolteae*, *Bifidobacterium dentium*, *Eggerthella lenta* and *Flavonifractor plautii* (also identified as a major core- influencer; **Fig 1B**) (enriched in >= 50% disease categories). As with the previously investigated properties, the 196 taxa could be placed into a graded spectrum based on the consistency of their health association patterns. This provided the basis for our third ranking index, which we referred to as the health-association score (HS).

Similar to stability scores, the health-association scores were computed using a boot- strapped iteration-based approach (**See Methods; Fig 3A; Supplementary Table 16**). Integrated examination of the three scores (Core-Influence:CS, Stability-Association:SS and Health-Association:HS) divided the 196 taxa into specific groups based on their association with three hallmark properties (**Fig 3C**). A core set of 18 taxa had all three association scores in the top 30% (**Supplementary** Figure 9; **Fig 3C**). We referred to these as the set of Health-Associated Core-Keystones (HACKs). Besides the 18 HACKs, some notable gut taxa like *Prevotella copri*, *Escherichia coli*, *Flavonifractor plautii*, *Collinsella aerofaciens*, *Gordonibacter pamaleae*, *Anaerostipes hadrus* and the two *Dorea* species, showed strong associations only with Core-Influence, but not with stability and health. Other taxa like *Paraprevotella xylaniphila*, *Ruminococcus lactaris*, certain *Bifidobacterium*, *Lactobacillus* spp. and *Butyrivibrio crossotus*, showed association only with health, but not with stability and influence. To account for these variabilities of association with the three individual properties, our next step was therefore to combine the three scores as a single combinatorial measure, called the ‘HACK-Index’.

### Final ranking of the gut microbial taxa based on a combinatorial HACK-index

The HACK index was calculated as product of two scores, namely the mean of the association scores of a taxon for all the three properties and a reward score that measures how similar (or evenly distributed) the three association scores were with respect to each other (**Fig 4A; Supplementary Table 17**). The top of this list was the well-recognized microbiome health marker *Faecalibacterium prausnitzii*, followed by *Bacteroides uniformis*. Other members in the top 20 on this list included multiple species of the *Roseburia*, *Alistipes*, and *Eubacterium* genera, and *Coprococcus catus*. Surprisingly, many taxa with high health-association (*Ruminococcus lactaris*, *Eubacterium eligens/ventriosum*) had noticeably lower HACK- indices because of their weaker association with either core-influence or stability.

**Figure 4:**
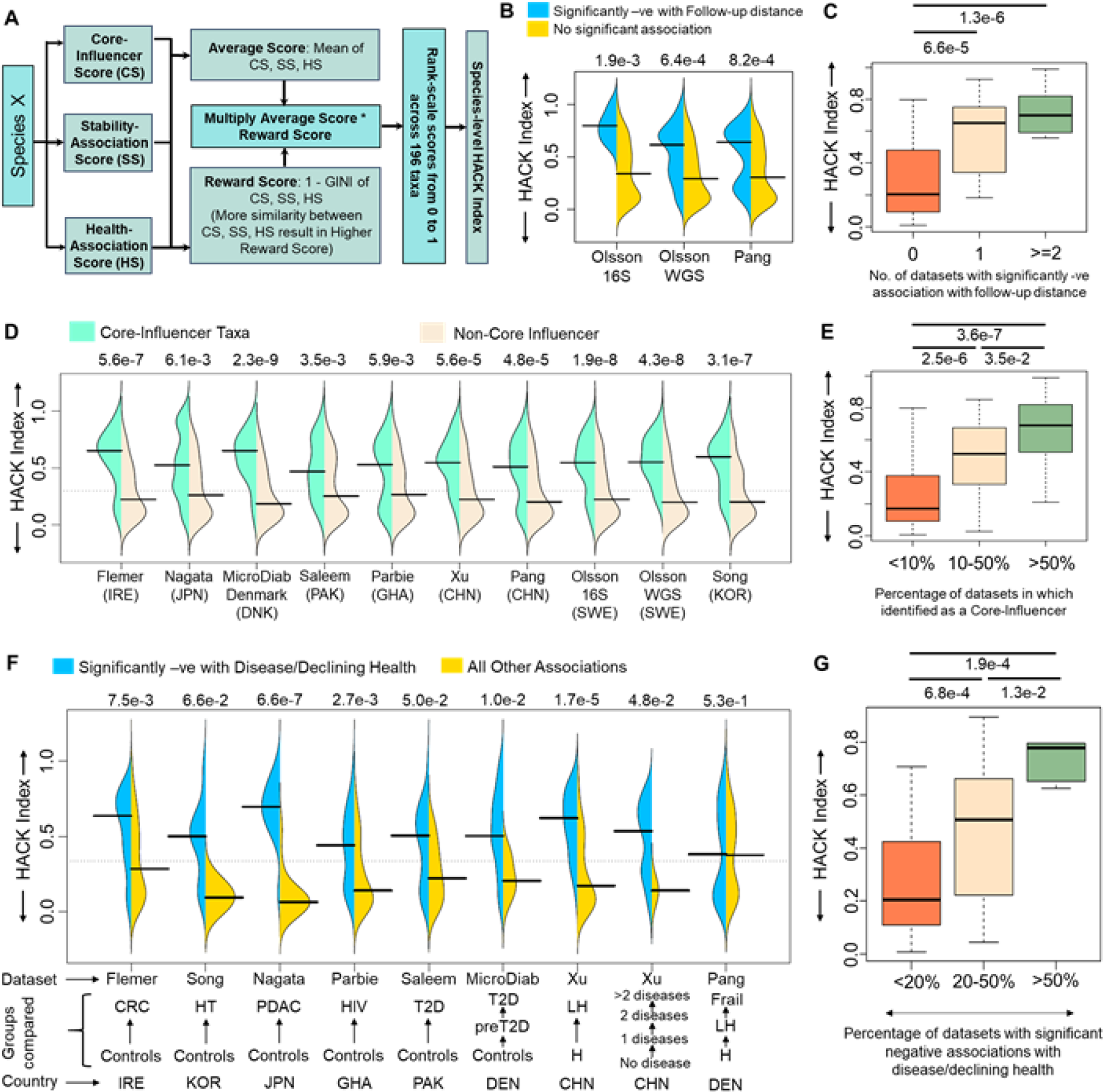
**Computation of HACK-index and the microbiome-level mHACK scores and the validation of the same in additional global validation cohorts**. **A.** Pictorially shows the approach used for computing species-specific HACK indices. **B.** Beanplots comparing the HACK-indices of stability-associated taxa (See Text) versus all other taxa across the three datasets. Stability-associated taxa had significantly higher HACK-indices as compared to others. **C.** Progressive increase in HACK indices in taxa-groups that are associated with stability with increasing consistency. The significant Mann-Whitney (MW) p-values of pairwise comparisons across groups are also indicated. Taxa detected with high consistency have significantly higher HACK-indices, followed by those identified with intermediate and low-consistency. **D.** Beanplot showing the significantly higher HACK indices of the core- influencers as compared to the non-core-influencer taxa identified for each of the cohorts. The MW P-values of the corresponding cohort-specific comparisons are also indicated. **E.** Progressive increase in HACK indices of taxa identified as Core-Influencers with low < intermediate < high consistency. The cross-group MW p-values are also indicated. **F.** Beanplots comparing the HACK-indices of taxa identified as negatively associated with disease/declining health (i.e. positive markers of health) with all other taxa in the nine comparative investigations performed in the validation cohort. For each beanplot, the dataset, the comparison groups and the nationality are also indicated, along with the corresponding MW p-values of comparison. **G.** Progressive increase in HACK indices of taxa that are identified to be associated with health with increasing degrees of consistency (lowest for taxa with the lowest frequency of health-association and highest for those identified with the highest consistency). The MW P-values of the corresponding comparisons are also indicated.

Interestingly, there were other unexpected and less-explored taxa on this list, namely *Fusicatenibacter saccharivorans*, *Agathobaculum butyriproducens*, and *Odoribacter splanchnicus* (at ranks 3-5), *Oscillibacter sp* (at 10), and two equol-producing taxa of the family *Eggerthellaceae* family *A.equilofaciens* and *A.celatus* (also identified as HACK- species; **Supplementary** Figure 10). The higher HACK index of these taxa as compared to other known markers like *Roseburia*, *Alistipes*, *Coprococcus* and *Eubacterium* was somewhat unexpected from the microbiome-health literature. Although all these taxa have previously shown sporadic positive associations with health (especially in cognitive function and healthy aging)^27–31^, the number of such studies are limited. These results indicated the applicability of the HACK-indices in identifying other promising members of the gut microbiome that could be explored for their beneficial effects.

The taxa with low HACK indices were dominated by oral taxa (*Actinomyces*, *Veillonella*, *Rothia* and *Streptococcus*) as well as by the known generally disease-associated lineages like *Ruminococcus gnavus*, pathobiont *Clostridium* (*C. ramosum*/*bolteae*/*clostridioforme*/*innocuum*,), *Eggerthella lenta*, *Klebsiella* and *Escherichia* (**Fig. 3C** and **Supplementary Table 17**). Notably, many of the earlier ‘beneficial’ taxa recently shown to have conflicting associations with health including *Akkermansia* and *Prevotella copri* (also shown to have high longitudinal variability)^32^, *Eubacterium hallii*, *Anaerostipes hadrus* and *Dorea longicatena*^9,33^, were identified with intermediate HACK indices. Thus, the obtained HACK-indices clearly reflect the known positive, negative as well as conflicting associations of specific taxa with health and microbiome stability.

### Validation of the HACK index in distinct globally diverse cohorts

To test the translational applicability of the HACK index, we investigated its reproducibility in 10 validation cohorts (**Supplementary Table 18; Supplementary Text 4**), that were not included in the 127 studies utilized for calculating the HACK-indices (**Supplementary Table 1**). These encompassed 4,400 gut microbiome samples, from industrialized populations in Europe (Sweden, Ireland, Denmark), East Asia (China, Japan and South Korea) to relatively non-industrialized regions like Pakistan and Ghana. The cohorts encompassed both cross-sectional and longitudinal samples from younger adults to centenarians, including matched patient and normal gut microbiomes from multiple diseases, namely T2D, Prediabetes, CRC, PDAC and HIV (**See Methods**). Within-cohort subsets comprising older subjects along with information pertaining to the health status of the individuals was also available. This cohort diversity allowed us to check the reproducibility of the HACK-index-based ordering of taxa across a wide range of geographies and diseases and measures of declining health.

We first investigated the association with microbiome stability. Three of the cohorts (‘Pang’, ‘Olsson_16S’ and ‘Olsson_WGS’) contained longitudinal samples, thereby enabling the validation of associations of HACK-index with longitudinal stability. For each cohort, we then identified the markers of stability as those taxa that showed significant negative association (Q<=0.15, Spearman Rho) with Bray-Curtis-follow-up distances (computed as in **Fig. 2A**). Across the three cohorts, the taxonomic markers of stability had significantly higher HACK-indices as compared to the non-markers (those showing non-significant or positive association with follow-up distances) (**Fig 4B**), thereby consequently reproducing the positive association of HACK indices with higher stability of the microbiome. Notably, the taxa identified as stability-markers in more than two datasets had the highest HACK-indices, followed by those detected as stability-markers in one dataset and significantly higher than those not detected as stability markers in any of the datasets (Mann-Whitney Test P=1.3e-6) (**Fig. 4C**). Thus, across each of the three cohorts, not only did the stability-markers have significantly higher HACK-indices compared to non-markers, but also the consistency with which they were identified as markers of stability significantly correlated with the computed HACK-indices.

We next investigated the core-influencer properties. Across each of the 11 cohorts, the Core-Influencer taxa (as in **Fig. 1B**) had significantly higher HACK-indices compared to the non-core-influencers (**Fig. 4D**). More importantly, the taxa having highest detection rates as Core-Influencers (detected in >50% cohorts) had significantly higher HACK-indices compared to those that were detected with medium (10-50% cohorts) and low (<10% cohorts) detection rates (Mann-Whitney P=3.4e-7 and P=3.6e-2, respectively), and those with medium detection rates as Core-Influencers had significantly higher HACK-indices as compared to those with the lowest (P=2.5e-6) (**Fig. 4e**). Thus, as observed for stability associations, not only was the association of HACK-indices with the core-influencers significantly replicated in the validation cohorts (including both industrialized and relatively-non-industrialized populations), the consistency of the identification as Core-Influencers significantly matched with their HACK- index-based ranking.

We then examined the reproducibility of the associations of HACK-indices with health. Five of the validation cohorts had microbiome data from patients with different diseases and matched controls (**Fig. 4F**). In addition, two of the cohorts also contained microbiome data from elderly subjects and centenarians that could be further stratified based on health-status as Healthy, Less-Healthy or Frail, as well as the incidence of 12 major diseases. This resulted in a diversity of comparative analyses. For each of the cohorts, we then computed the association of the abundance of the different taxa with disease using Robust Linear Regression models (**See Methods; Supplementary Text 4**). In eight of out the nine comparisons tested, the taxa having significant negative association with disease/declining health status (i.e. the markers of health) had lower HACK indices as compared to those not showing such associations (significantly in seven of the cases). Furthermore, the consistency of the association of the different taxa with health in this targeted analysis strongly correlated with their previously computed HACK-indices (**Fig. 4G**).

Thus, our validation investigations showed the HACK-based ranking order of the taxa is significantly replicated in terms of the association of the individual taxa with respect to all three properties of core-influence, stability and health.

### Links between HACK-indices and responses to specific microbiome-associated therapeutic interventions

As a further test of their translational applicability, we next examined the links between the HACK-index based taxa ordering and different microbiome-associated therapeutic interventions. Here, we investigated whether this ordering/ranking also correlated with the taxon-specific associations and/or responses with consumption of (specific constituents of) therapeutic (healthy) dietary regimes (e.g. Mediterranean diet or MED-diet) and also with respect to their modulatory effect on the host-response to specific therapies (e.g. Immuno- checkpoint therapy (ICT) in cancer patients (**Supplementary Text 5**).

Here, we first studied data from a MED-Diet intervention trial previously investigated by Meslier *et al*^34^ (62 subjects/individuals, 30 with MED-diet intervention and 32 with habitual-/CONTROL-diet, with gut microbiome and diet data profiled at three time-points (baseline, 4-weeks and 8-weeks post-intervention). Across 4 of the 6 time-point-subject-group combinations, we observed significant positive relationships between taxa-specific HACK indices and the MED-diet compliance scores of the corresponding subjects (**Supplementary** Figure 11A). Thus, the higher the HACK-index of a taxa, the more positive was its abundance correlation with increasing adherence to the MED-diet. This was further evident while investigating taxa abundance associations with respect to the consumption of individual dietary components. HACK indices showed a significant positive link with the consumption of MED- associated dietary components like fruits/vegetables, dietary fibers and vegetable proteins (i.e. the higher the HACK index of a taxa the more positively was its abundance associated with the consumption of these items) and; a negative link with consumption of lipids, saturated fats and animal proteins (components reduced during MED-diet) (**Supplementary** Figure 11B).

Notably, the Week4-baseline abundance-differences of the 18 HACK species (**Supplementary** Figure 9) not only showed significantly different patterns between the CONTROL- and the MED-diet groups (PERMANOVA R-squared: 0.05; P=0.016; **top panel Fig. 5A**), but this alteration pattern significantly correlated with diet linked Week4-baseline dietary alterations (PROCUSTE: R=0.28; P=0.001). No such pattern was observed for the other taxa. 70% of subjects in the MED-diet group had an increase in the cumulative abundance of the HACK-species between the two time-points (the other taxa showed the opposite pattern: Fishers’ exact test: P=0.019) (**Fig. 5B**). Furthermore, the abundance alterations of 8 (of the 18) HACKs correlated positively with dietary changes in the MED-diet group (only 3 associated with changes in the CONTROL group) (**Supplementary Text 5; Fig. 5C**). Thus, the high HACK-index taxa are positively enriched by increased adherence to the MED-diet.

**Figure 5.**
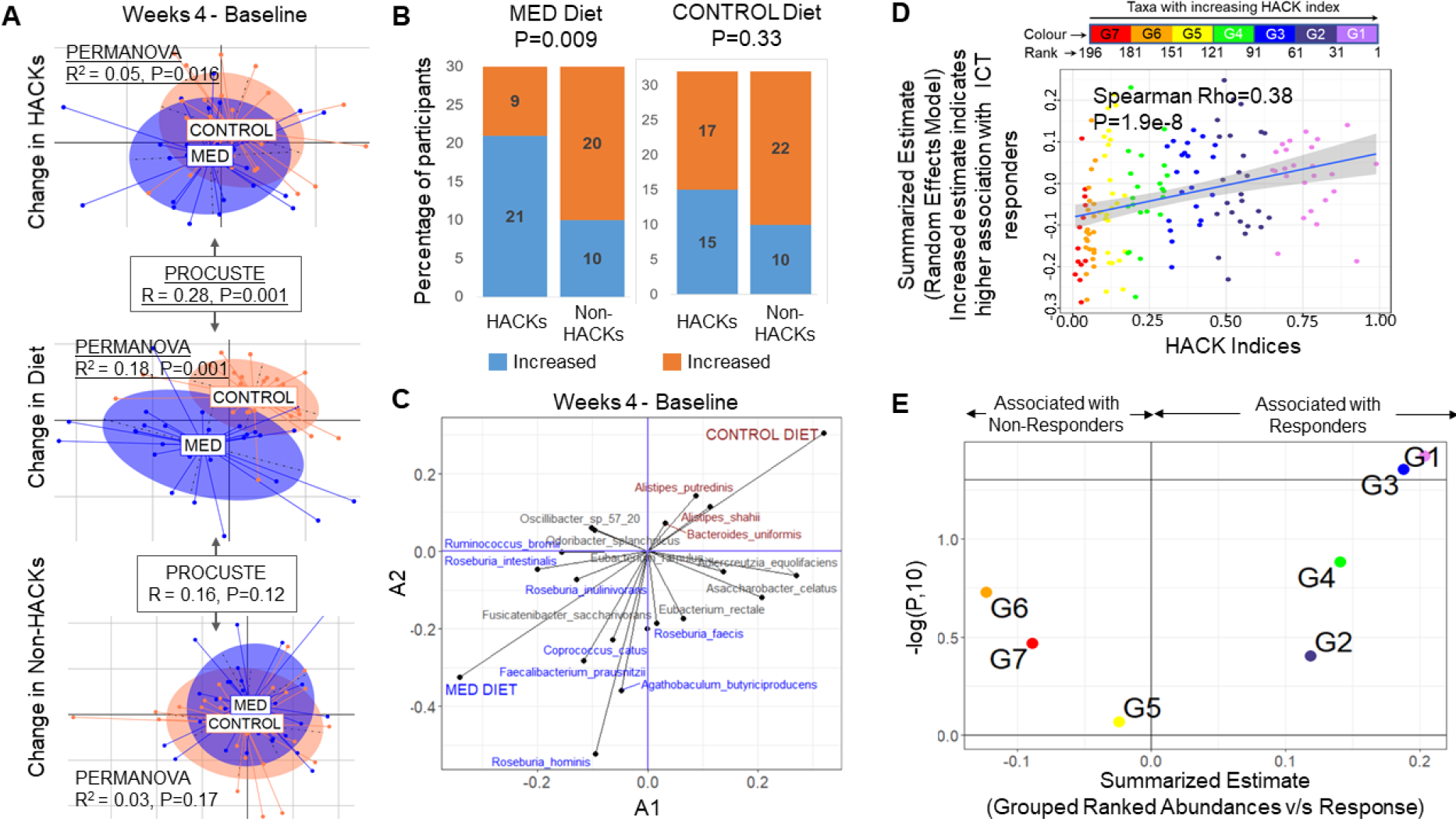
**Association of High HACK index with positive (or favorable) response to Microbiome-associated therapeutic interventions**. A. Principal Coordinate Analysis (PCoA) (of Meslier *et al* MED-diet cohort data) performed on the Weeks4 – baseline differences of: ranked abundances of the 18 HACKs (Supplementary Figure 9) (top panel); the ranked intake frequencies of the different dietary components (middle panel) and; the ranked abundances of the non-HACK (all except the 18 HACKs) (bottom panel). Subjects (points) from CONTROL-diet and MED-diet groups are shown in different colours. PERMANOVA results to compute the group-specific significant differences in the alteration patterns are also indicated, along with PROCUSTE-analysis results to judge the significance of the correspondence between dietary alterations and the different microbiome alteration profiles. **B.** Barplots showing that the MED-diet group has a significantly higher number of subjects with increased ranked abundance of HACKs (as compared to non-HACKs). No such pattern is observed for the CONTROL-diet group. **C**. Scatterplot where in each point indicates a HACK taxon with the x- and y-coordinates being correlations of its abundance change with component1 and component2 of the diet-change PCoA, respectively. The centroids of the diet-change PCoAs for the MED-diet and CONTROL-diet group are also shown. Those showing positive correlation with diet change in MED-diet group are shown in blue and those showing negative are in red. **D.** Shows the results of a Random Effects Model Meta-Analysis, where taxa with increasing HACK indices tend to have more positive summarized estimates (indicating responder association). The corresponding positive correlation along with its significance is also indicated. To further elaborate this, taxa were divided into seven different consortia (G1- G7) based on decreasing ranks of HACK-indices and coloured differently as explained in the top panel. **E.** Volcano-plot showing the results of the meta-analysis investigating the association of the cumulated ranked abundance of the seven taxa consortia (mentioned above; top panel of D), with ICT response across the five cohorts. G1 consortia containing the top 30 taxa in terms of their HACK index shows the strongest association with responders.

Next, we investigated the links between the HACK indices and the gut microbial determinants of host response to immune-checkpoint therapy (ICT) in cancer patients. The baseline gut microbiome composition (e.g enrichment of *Akkermansia muciniphila*, *Roseburia* in responders) has been associated with ICT responses, but the exact determinants have been highly cohort-specific showing limited replicability/translatability across cohorts^35^. Here, we investigated five such cohorts to test whether, despite cohort-specific variabilities, we could still identify taxa groups (or consortia), showing reasonable across cohorts and if these groups were correlated with our HACK-index-based taxa order (**Supplementary Table 19; Supplementary Text 5**). Taxa-specific random-effect model based meta-analysis validated the previously reported positive associations of *A.muciniphila* and *Roseburia* with ICT response (**Supplementary** Figure 11A). In addition, the analysis revealed a significant positive relationship between the extent of association of different taxa with responders (measured as summarized estimates of the random-effect models; positive estimates indicating association with responders) and their HACK-indices (Spearman Rho = 0.39; P-value=1.9e-5) (**Fig. 5D**). The pattern was also in the study-specific consistencies of these associations (**Supplementary** Figure 11B). To further probe this positive link, we divided the 196 taxa into groups (or consortia) based on HACK indices (**Fig. 5D**; top panel) and repeated the meta-analysis. The G1 (the taxa-group containing the top HACK-indices) showed a significant positive association with ICT responders across the five cohorts (along with G3). None of the other groups showed any significant association (**Fig. 5E**). Thus, the top taxa in the HACK-index order (as a consortia) are significantly associated with positive responses to ICT. Our next objective was to identify the functional determinants contributing to the HACK-indices and particularly the functional signatures of taxa with higher HACK-indices.

### Identification of functional signatures associated with HACK-indices

We first examined the experimentally-validated species-level metabolic functionalities independently collated from published studies^11,12^, and used in previous studies^4,13^. (**See Methods; Supplementary Table 20**). Using a two-step approach combining Random Forest and logistic regression models, we identified 22 metabolic functionalities showing strong association (p-value < 0.05) and 15 metabolic functionalities having marginal association (0.05 <= p-value < 0.1) with taxa-level HACK-indices (**See Methods; Fig. 6A-B; Supplementary Table 21**). The RF models, built using the taxa-specific detection profiles of these 22 top metabolic functionalities, yielded a significant positive correlation between the actual and the predicted HACK-indices (obtained using the RF-prediction models built upon the) (Spearman Rho = 0.52, P-value = 1.3e-9) (**Fig 6C**), indicating that HACK-indices have strongly associated metabolic signatures.

**Figure 6.**
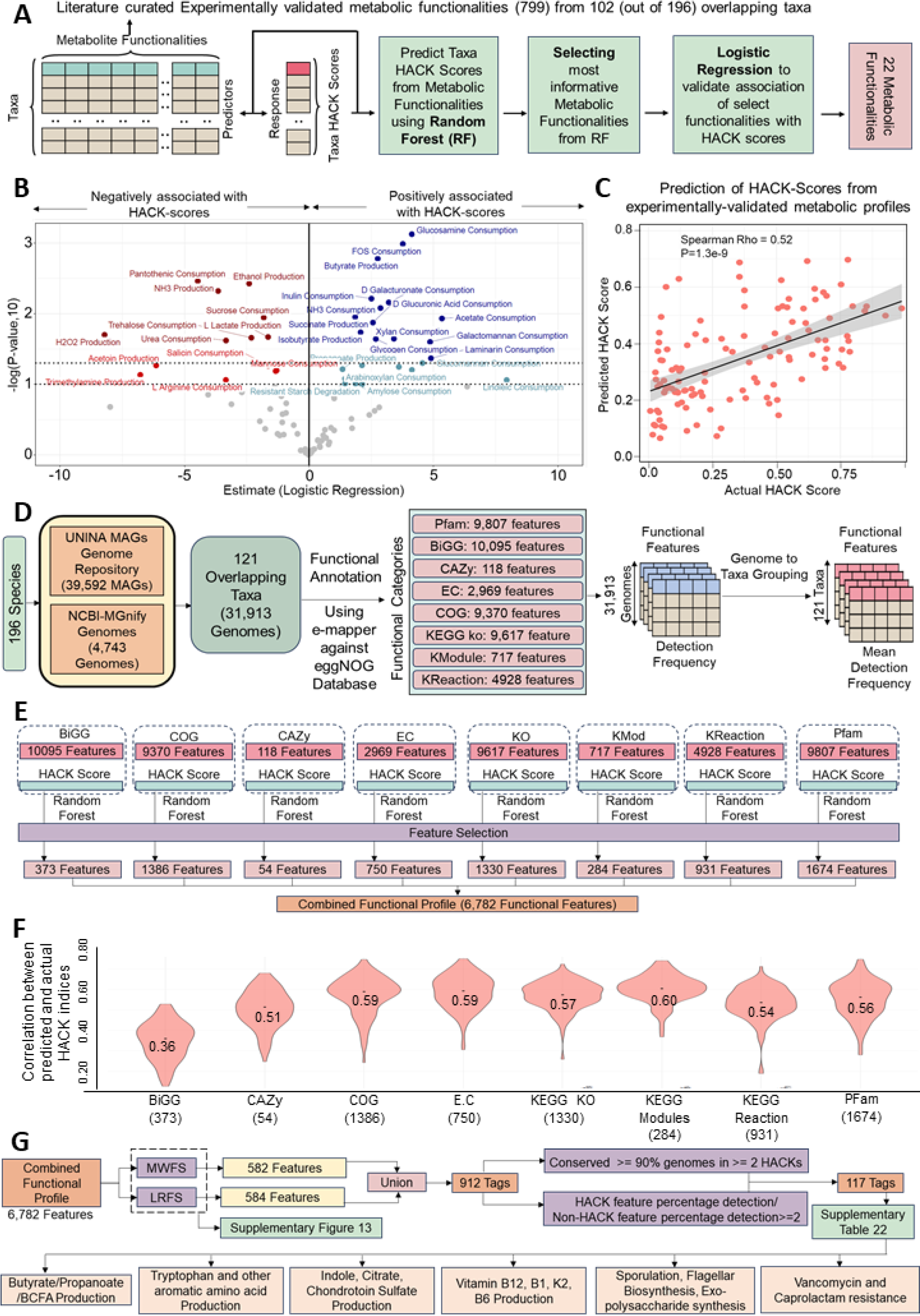
**Identification of Functional Signatures associated with HACK indices and the HACKs**. **A.** This panel shows the approach used for identifying the experimentally validated metabolic functionalities that are predictive with the taxa-specific HACK-indices. **B.** Graphing of the metabolic functionalities (identified in A) based on their directionality of the association with HACK indices. Metabolic functionalities positively associated with HACK indices are highlighted in either blue (P <= 0.05 post logistic regression (LR)) or cyan (0.05 < P <= 0.1 post LR) and those that are negatively associated highlighted in either red (P <= 0.05 post LR) or light orange (0.05 < P <= 0.1 post LR). **C.** The out-of-the-box (OOB) correlations between Actual and Predicted HACK-indices obtained using Random Forest (RF) models built only using the features shown in **B. D.** Schematic representation of the functional feature profiling strategy adopted for 31,913 genomes obtained from the 121 taxa (overlapping with our list of 196 taxa) to obtain their species-level functional detection profiles using the eight different schema. **E.** The approach used for obtaining a select list of features from each schema (encompassing a total of 6782 features). **F.** Violin plots showing the distribution of correlations between the actual and predicted HACK-indices obtained for taxa across 50 iterations of RF models (2-Fold Cross Validation: Trained on 50% taxa and tested on 50% taxa), using the select features identified for each functional schema. The median correlations for each functional schema are also indicated. **G.** Approach used to further reduce the set of 6782 HACK index-associated functional features to identify a select list of 117 functional features significantly enriched in only the HACK taxa.

Taxa with high HACK-indices showed significantly positive association with butyrate, isobutyrate and succinate production and marginally positive association with propanoate (**Fig 6C**), indicating that taxa with higher HACK-indices tended to be enriched with genes for production of major short chain and branched chain fatty acids. Species with higher HACK- indices also had unique nutrient preferences characterized by specific complex carbohydrate consumption, like fructo-oligosaccharides (FOS), xylan, galactomannan, xyloglucan, glucosamine, glucuronic acid and marginal enrichments in the consumption of glucomannan. Taxa with lower (or decreasing) indices were characterized by the production of many metabolites with detrimental effects like hydrogen peroxide (H2O2), ethanol, trimethyalmine (TMA) and ammonia (NH3) and importantly with the nutrient preferences linked to consumption of several simple sugars like Sucrose and Fructose. Most TMA-producing gut-associated taxa including pathobiont Clostridia (*C. hathewayi/citroniae/bolteae/asparagiforme/clostridioforme*), *Escherichia* and *Klebsiella* species were identified with HACK indices in the lower tertile (**Supplementary Table 17**). We previously observed a similar functional signature associated with a transition to a non-communicable disease-associated microbiome in our previous studies on Elderly Irish and the Irish Travellers^14,15^. By validating the connection between previously known experimentally validated metabolic functionalities with HACK-indices, this association analysis clearly provided the proof-of-principle of utilizing HACK-scores as a measure of identifying functionalities associated with generic microbiome health and disease. However, the number of associated metabolic functionalities was limited and the level of significance of association for a majority of these was also low. This was primarily because not all metabolic functionalities were experimentally validated across all species-level taxa and not all species-level taxa had information pertaining to their experimentally validated metabolic profiles.

We extended this investigation to genome-level functional annotations using eight different schema for each of these taxa and attempted to identify a comprehensive list of functional profiles significantly associated with HACK-index and the HACK-species (**Summarized in Fig. 6D-G and Methods; Supplementary** Figure 12). This investigation, explained in detail in **Supplementary Text 6**, identified a select set of 912 functional features (or tags) as the collective output from the 8 schema (encompassing a total 47,621 tags). From these, we selected a core-list of 117 functional tags that were significantly enriched the 18 HACK taxa (**Supplementary Table 22**). These 117 functional tags represent a wide range of functionalities, ranging from (anti-inflammatory) butyrate/propanoate production, production of multiple vitamins, neuro-active amino-acids like tryptophan as well as their anti-inflammatory beneficial derivatives like indole/IPA, chondroitin sulfates as well as a range of diverse functionalities that are hitherto unexplored. These represent candidates that can be investigated further from an evolutionary, mechanistic and pre-clinical perspective to unravel the key functions that characterize the health-associated keystone members of the human gut microbiome and aid in the identification and formulation of novel-biotherapeutics.

## Discussion

While the diagnostic and prognostic value of an index of microbiome health seems self-evident, it may also help identify taxonomic targets for microbial therapeutics. Prioritization of taxonomic targets for restoration therapies requires the evaluation of gut microbial members based on their association with host health and also the stability of the gut microbial ecosystem. Our approach proposes a gradient or continuum of 196 universally-prevalent species-level taxa ranked for their association with three specific criteria of host health as well as that of the gut microbiome ecosystem. By using a global discovery dataset of ∼40,000 adult gut microbiomes (both cross-sectional and longitudinal) from 42 countries and including 28 different disease conditions, we found that not all markers of host health are necessarily prevalent keystone members (influential members associated with microbiome stability) associated with ‘normal’ microbiome health and not all keystones are necessarily associated with positive host health. Recent studies have shown that while some probiotic candidates e.g. *Bifidobacterium*, show low intra-individual variability, others like *Lactobacilli*, are associated with high temporal variability (indicating their association with an unstable microbiome state)^32^. This is also observed in our ranking order computed in this study (**Supplementary Table 17**).

By incorporating data from different continents with diverse life-styles, we aimed to mitigate the variability of the normal microbiome across different populations and ensure that the HACK indices are universal. Consequently, many of the species previously observed to show region-specific associations with normal (or apparently healthy) gut microbiomes, e.g. *Prevotella copri* in the non-industrialized and multiple Bacteroides species (except *B. uniformis*) with industrialized populations, are identified with lower HACK indices. By incorporating gut microbiome data from patients of multiple diseases, we are also able to discriminate taxa, showing context-dependent association with health (e.g. *Akkermansia* and *Dorea*).

Nevertheless, there is a clear correlation with respect to all three hallmark properties of health and microbiome-resilience: prevalence/community-influence in non-diseased subjects, longitudinal-stability and host health, which is captured by the HACK-indices. This rank order is led by the anti-inflammatory and candidate next-generation probiotic *Faecalibacterium prausnitzii,* and followed by several less-explored members of the gut microbiome such as *Fusicatenibacter saccharivorans*, *Odoribacter splanchnicus*, *Agathobaculum butyriproducens*, *Adlercruetzia equilofaciens* and *Asaccharobacter celatus*. The results suggest a role for these taxa as potential as biotherapeutics.

The reproducibility of this index as a single indicator of core-influence, stability and health is further shown in 10 geographically distinct validation cohorts encompassing more than 4,000 cross-sectional as well as longitudinally sampled gut microbiomes (additional to the 127 cohorts considered for computing the indices). A crucial application that is observed for the HACK indices is for identification of gut microbial drivers of a positive response in two diverse microbiome-linked therapeutic interventions (MedDiet and ICI Therapy in Cancer). For both the interventions, one of the gut microbial features most consistently associated with a positive response is an enrichment of taxa with the highest HACK indices (the top 18 HACK taxa for MedDiet and the consortia of taxa with the top 30 HACK indices). This supports the use of the index to select consortia of taxa for use as synergistic therapies.

Finally, our investigation identified specific functional/metabolic signatures associated with the HACK indices and 18 HACK taxa. Functionalities like SCFA production, consumption (nutritional preference) for prebiotics such as fructo-oligosaccharides, inulin, xylan, galactomannan, production of tryptophan, indoles, chondroitin sulfate and multiple vitamins, along with transmembrane proteins with signal transduction functions indicate a range of functionalities encoded by these taxa that can impact upon diverse aspects of host-physiology and host-microbiome communications, including cell-to-cell interaction, gut-brain axis and immune modulation, which need further experimental exploration.

It is noteworthy that the HACK indices apply to adult human gut microbiomes but the ranking order is likely to vary in infants and adolescents. At present, we are extending our analyses to infants and the elderly. With a further increase in publicly available gut microbiome data, these HACK indices may undergo further refinements, including incorporation of more datasets from non-industrialized and rural populations. In the present study, we noticed a lack of longitudinally sampled datasets from less industrialized populations. Despite this limitation, the HACK indices were computed by an investigation of greater than 45,000 well annotated adult gut microbiome datasets from 144 cohorts representing all six major continents, making it one of the most comprehensive investigations to date, to our knowledge.

The HACK indices represent a step toward rational prioritization of gut microbial species for exploration as potential microbiome-directed therapeutics. Moreover, previously unexplored HACK-associated functionalities may help identify pathways and metabolic capabilities associated with general microbiome health and stability.

## Methods

### Collation of gut microbiome datasets

The discovery phase of this study involved three systematic global investigations of grading the various gut microbial taxa based on their associations with three key hallmark properties of microbiome health and resilience. Overall, the investigation encompassed 39,926 gut microbiomes from adult individuals (age >= 18 years) collated from 127 study cohorts covering 42 countries and six continents.

The details of the 127 study cohorts are provided in Supplementary Table 1 and described in Supplementary Text 1.

### Data Processing and Generation of Taxonomic Abundance Profiles

For the datasets downloaded from the curatedMetagenomicData repository, the metaphlan3 generated taxonomic profiles along with the sample metadata were already available^20^. For the rest of the whole genome sequencing datasets, the same protocol as adopted by the curatedMetagenomicData was utilized. All 16S derived gut microbiome datasets were downloaded from the corresponding data sources (as denoted by the accession number columns in Table) and processed using the SPINGO taxonomic classification tool^36^. The corresponding metadata for all the datasets that were not part of the curatedMetagenomicData, were obtained by looking into the respective research papers. We thank the authors of the Arivale cohort for providing us with the metadata of the corresponding participants.

### Rank-scaling of values

A key component of the computation of the different association scores was the rank-scaling function utilized here. For a given array of values ‘x’ of length k, denoted by x = {x1, x2, x3, x4, …., xk}, the values are first ranked. Next, the corresponding rank-scaled values, denoted by 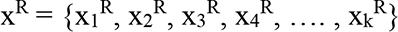, of this set were obtained as:

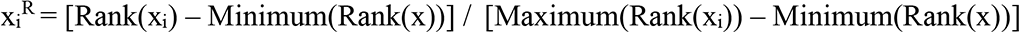

The transformation was essentially an approach of ranking the values and then converting the ranks to values between 0 and 1.

### Computation of the Core-Influencer scores

The Core-Influencer scores for the different taxa were computed simply by counting the number of the cohorts a given taxa was identified as a Core-Keystone and subsequently rank-scaling these numbers obtained for all the taxa (as described above) to values from 0 and 1.

*Identification of Core-Keystones for a given cohort*: For a given study cohort ‘Y’, a Core- Influencer was identified as a taxon that satisfied the following two criteria:

- Prevalence of >= 70% (i.e. detected or present in at least 70% of the samples or gut microbiomes of Y)
- Community-Influence Rank >= 70%. The procedure to compute the Community- Influence Rank as well as the approach used to select the threshold of 70% is described below.

For each taxa, the absolute Community-Influence was first calculated using the Remove- Renormalize-Relate approach (3R) as described below.

*Computation of community influence using the Remove-Renormalize-Relate (3R) approach*: The Remove-Renormalize-Relate (3R) approach is described schematically in **Fig 1B**. Briefly, for a given study cohort A, to measure the community influence of a given taxon (X), we first removed the abundances of X from all microbiomes belonging to cohort and recomputed the abundance profiles of the microbiome by renormalizing the abundances of the remaining taxa. The abundances of all remaining taxa were then renormalized and the community variations of the resulting microbiome profiles were then computed using Principal Coordinate Analysis using the Bray-Curtis distance matrix. The PCoA of the 3R approach was performed using ‘dudi.pco’ function of the ade4 package v1.7.22 of the R-programming framework. The input was the Bray-Curtis Distance matrix containing the distances between each microbiome-pair provided as the input. The Bray-Curtis distances were computed using the ‘vegdist’ function of the vegan R package v2.6.4.

The removal of the test taxon X during the computation of community-wide variations ensured that the estimated variations in community composition were not simply due to the large variations in the relative abundances of X, but were instead solely based on the differences in the mutual representation of remaining microbiome members. The community-influence of X was then computed by inter-relating the abundance of X with the community-wide variations calculated (after the ‘removal’ of X) as described above. The envfit approach, encoded in the function with the same name in the vegan package as described above, was utilized for this purpose. The method provides two outputs: an R-squared value, indicating the extent of community- influence the abundance of X has on the ordination scores (depicting the absolute effect the abundance of X has on the community-wide variations of the remaining taxa); a p-value is the probability of observing the R-squared value by random chance (being computed using a permutation test) and indicative of the significance of the effect. Taxa were then rank-scaled (as described above) based on their community-influence (R-squared value obtained using envfit) (**Fig 1B**).

The 3R approach is derived from on the recently published Empirical Presence-Absence Interrelation (EPI) approach^23^, with exception that it is based on abundance variation rather than the EPI’s presence-absence information. We did not adopt the presence-absence based approach as the core-influencers are expected to be prevalent, classifying microbiomes simply based on the presence-absence of these members may result in a bias in the number of microbiomes belonging to different groups. Furthermore, a taxon is also likely to have variations in its community-influence depending on how abundant is it in a microbiome.

### Association of Core-Influencer Scores with study-specific microbiome and host properties

*Abundance*: In this investigation, we checked and ensured that the detection as Core- Influencers was not a simple reflection of abundances of the identified species (i.e. only the highly abundant species being identified as Core-Influencers). For this purpose, we computed the study-specific mean abundance of each species and correlated the same with their study-specific community-influence (rank-scaled R-squared of the envfit) and prevalence using Pearson correlations.

*Cohort Life-Style and Cohort Sequencing-Type:* All gut microbiomes comprising the study cohorts were derived from individuals in the age-range >= 18 years. Individual microbiomes had a mix of different host demographics like age, gender, BMI etc. Some of the studies had a mix of nationalities. However, there were certain properties that were especially cohort-specific and could be used to broadly categorize almost all cohorts. These were the microbiome profiling strategy (referred to here as ‘SequencingType’) (amplicon-based or 16S-based; whole genome sequencing or WGS-based) and the life-style of the study populations (referred to here as ‘CohortLifeStyle’). For the later, we checked the individual research papers pertaining to each study and subsequently categorized the life-style pattern of the study populations into three groups, namely Industrialized-Urban, Urban-Rural-Mixed and Rural-Tribal (**Table 1A**). As the names suggest, the first category was “IndustrializedUrban” (consisting of studies from Europe, North America and East Asia, where a majority of individuals reside in Industrialized Societies). The exceptions to these were the cohorts of Irish Traveller from Ireland (KeohaneDM 2020) and a large Chinese population cohort (He 2018), which have investigated individuals from mixed urban/settled and rural/nomadic life-styles^6,37^. These two studies, along with those focusing on populations from India, Fiji, Native New Zealanders, Pacific Islanders, and other South American non-nomadic cohorts, having a mixture of both Urban and Rural sub-cohorts, were placed into a second category called ‘UrbanRuralMixed’. The third category consisted of nomadic/semi-nomadic or settled-rural cohorts from Africa, Peru and Mongolia and was tagged as ‘RuralTribal’. Investigating with the SequencingType and CohortLifeStyle factors on the properties (prevalence, community-influence) of the gut microbiome members as well as the detection pattern of Core-Keystone taxa across cohorts was performed as described below.

We first performed study-level PCoAs (Euclidean distance for Prevalence and Community-Influence Rank; Jaccard Distance for Core-Keystone Detection). Subsequently, for each property, pairwise distances were computed between the studies by computing the Euclidean distances between the corresponding Principal Coordinates. Subsequently, by using the Permutational Multi-Variate Analysis of Variance (PERMANOVA) approach, we investigated if these observed variations between studies could be explained by their ‘SequencingType’ or the ‘CohortLifeStyle’ of the study populations. PERMANOVA was performed using the adonis2 function of the vegan R package (described above).

### Computation of Stability-Association scores by investigating a distinct set of longitudinal cohorts from diverse geographies

*Calculation of Follow-up Distances*: The key methodology corresponding to this specific part of our investigation was the sample-specific Follow-up distances. As described in the Results, sample-specific ‘Follow-up Distance’ (D^x^t) was calculated as the distance between the species-level microbiome composition of a given time-point sample S^x^t (from a given subject ‘x’ at time-point ‘t’) with that collected from the same subject ‘X’ at the immediate subsequent time- point S^x^t+1. Across the 23 different studies, this distance was calculated for each of the 7,134 gut microbiome samples with follow-up time-points (from all the 2,234 subjects across the 23 studies). For those samples of a given subject which did not have a subsequent time-point, the Follow-up distance was assigned as ‘NA’. The Follow-up distances were also computed using two different distance measures, namely Aitchison and Bray-Curtis (described below):

*A. Aitchison-based Follow-up Distance*: Here, we first performed the Centered-Log Ratio (CLR) Transformation of the abundance values. To cope with 0 values in abundance profiles, prior to CLR-transformation, the normalized species-level relative abundance profiles were initially transformed by adding a pseudo-abundance value of 0.0001 to all the abundance values (‘x + 0.0001’). The CLR-transformation was subsequently performed on these abundance profiles, using the ‘clr’ function of the compositions package in R v2.0.4. After the CLR-transformed abundances were obtained, to ensure that all the minimum CLR-transformed values (corresponding to the 0 values in the abundance profiles) are the same, we performed a second transformation of the CLR-transformed abundances using the formula ‘x-min(x)’ where ‘x’ is the species abundance and ‘min(x)’ is the minimum of the species abundance in a given sample. After the generation of CLR-transformed abundances, the Follow-up distance between the microbiome composition of a given sample S^x^t and S^x^t+1 by computing the Euclidean distances between the CLR-transformed abundances of same species across the two samples. For this, we used the vegdist function of the vegan R package setting the ‘method’ argument of this function as ‘euclidean’.
*B. Bray-Curtis Follow-up Distance*: The Bray-Curtis Follow-up distance between a given microbiome composition S^x^t and its subsequent follow-up microbiome S^x^t+1 was computed by providing both the abundance profiles as a matrix to the same vegdist function, but this time setting the ‘method’ argument as ‘bray’.

*Quantifying the association of the different taxa with longitudinal stability of the gut microbiome using boot-strapped iterations*: The follow-up distances are an inverse measure of stability (higher distance indicates higher variation across time-points and lower microbiome stability). Thus, to investigate this, on the gut microbiomes belonging to the 23 longitudinal study cohorts, we performed 10 iterations each time selecting 12 of the 23 study cohorts. For each iteration, we computed an iteration-specific score for each taxa by quantifying the stability-association of the different taxa can thus be quantified by associating their abundances in a gut microbiome at a given time-point with the corresponding longitudinal variation of the same microbiome at the subsequent time-point (measured by the follow-up distance) as described below.

In each iteration, we performed meta-analysis using Random Effect models to compute the association of the different taxa with both the Aitchison and Bray-Curtis distances across the 12 study cohorts. For this purpose, first, within each cohort, we considered all those microbiome samples with follow-ups (i.e. samples collected from the same subject at a next subsequent time-point). For every study cohort, the Follow-up distance of the microbiome composition of each sample S^x^t with that of its immediate follow-up sample S^x^t+1 was calculated as described above. Subsequently, for a given taxa ‘Y’, the association of its abundance across each such sample S^x^t with the corresponding obtained follow-up distances D^x^t was then evaluated using robust linear regression models (RLMs). The RLMs were computed using the rlm function of the MASS R package v7.3.58.3. The RLMs were computed for both the Bray-Curtis and Aitchison Follow-up Distances. The p-values (of significance) of the RLMs were computed using the robust F-test (f.robftest) function of the sfsmisc R package 1.1.15. For a given taxa ‘Y’ and a given measure of Follow-up Distance (Aitchison or Bray-Curtis), the estimates of the RLM for a given study was obtained and treated as the effect-size of the association for that study. For a meta-analytic investigation across studies, the effect-sizes between the same taxa-Follow-up-distance pair obtained for each of the 12 longitudinal studies were then provided as inputs to a Random Effects Model-based (REM-based) Meta-Analysis. This was performed using the ‘escalc’ and the ‘rma’ function of the ‘metafor’ R package version 4.2.0.

For each of the 196 taxa, the summarized estimates, the p-value of the Random Effects model was obtained. The set of p-values obtained for all the taxa were then corrected to the Q-values using the Benjamini-Hochberg correction, performed using p.adjust function of R base package with method as ‘fdr’. Additionally, we also computed another property of the association called consistency, which referred to percentage of study-cohorts where the directionality of the association was the same was the same as the summarized effect size. The consistency gave an indication of how robustly the association captured by the meta-analysis was also reflected within the individual study cohorts. Finally, an iteration-specific (say given iteration ‘i’) stability-association score for a taxon ‘j’ was computed as:

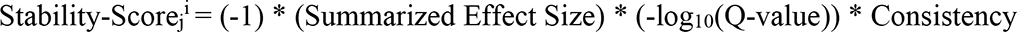

We obtained separate Stability-Association scores with respect to the Aitchison and Bray-Curtis follow-up distances, and then computed the mean of the same. The mean of the Aitchison and Bray-Curtis stability-association scores for all 196 taxa obtained in iteration ‘i’ were then rank-scaled from 0 to 1 to obtain the iteration-specific mean-rank-scaled stability- association scores. The mean-rank-scaled stability-association scores across all 10 iterations were then averaged to obtain the final overall Stability-Association score. The iterative approach ensured that the association scores thus obtained were robust to variations and not biased by the associations of a given taxa with certain datasets of high sample sizes. The multiplication of the Summarized Effect Size by (-1) was performed because we were computing Stability-Association Scores and the Summarized Effect Sizes were computed with respect to the follow-up distances (which was an inverse indicator of Stability).

*Computation of taxa-stability-association scores after adjusting for abundance for the specific taxa*: The scores were computed in exactly the manner as described above. The only modification was that within the cohort-specific Robust Linear Regression models associating the abundance of a given taxon ‘j’ with a given (Aitchison or Bray-Curtis) follow-up distance, we added the abundance of ‘j’ across the time-points as a covariate:

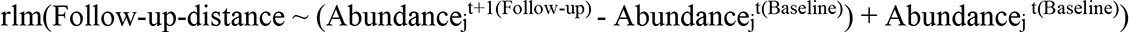

This investigation validated whether the identified associations with Follow-up distances (or Microbiome stability) are not simply driven by the across-time-point abundance variations of a taxon itself that affects the Follow-up distances.

*Computation of taxa-stability-association scores within different subject groups*: Another key confounding factor to investigate here was the phenotypic category of the study subjects. The 23 study cohorts considered herein contained longitudinal microbiome samples collected from apparently non-diseased controls, Ulcerative Colitis and Crohns’ Disease patients, Fiber intervention trial participants (with all of the four categories being sampled as part of multiple studies (and consequently grouped into the corresponding sub-groups), along with Irritable Bowel Syndrome and Chronic Liver Disease patients, antibiotic-treated, drug-resistant Enterobacteriaceae harbouring subjects and Fecal microbiome transplation patients (all grouped into ‘Others’ category) (**Table 1B**). To ensure that the observed pattern of associations is not confounded or biased towards specific subject phenotypes constituting the study cohorts, we also repeated the above examinations for each phenotypic sub-group, each time considering the subjects belonging only to that sub-group. The reproducibility of the overall association patterns individually within the different phenotypic group was then investigated.

### Computation of Health-Association scores using global cohort of matched disease-control subjects covering 28 different diseased conditions

As for the previous steps, we specifically investigated the same 196 species as used for the computation of the Core-Influencer and Stability-Association scores. The selection of the study cohorts and the 28 disease conditions have been described previously in this methods section. As in the previous section, the Health-Association scores were computed using a similar bootstrapped iterative approach as adopted for the Stability-Association scores. Here also, we performed 10 iterations, each time selecting 18 (of the 28) disease/clinical conditions (**Fig 3A**).

Within each iteration for each disease and a given taxon, we determined whether the taxon was associated with health or disease by comparing its abundance in control samples with respect to the diseased. For a given disease, we combined all study cohorts corresponding to the disease (based on the criteria described above) (listed in **Supplementary Tables 12-14**). The abundance comparison was performed using a two-step approach. First, we checked the directionality of the change (i.e. whether the bacteria is more abundant in disease or in control-matched samples). Directionality was computed as either 1 (Mean Abundance in Disease > Mean Abundance in Controls) or -1 (Mean Abundance in Controls > Mean Abundance in Disease) or 0 (No Difference). Next, we checked what is the significance of the difference in the abundance of the species between control and disease samples. For this purpose, we performed Mann-Whitney test using the disease and matched-control samples. The p-values were obtained and were also subsequently corrected using the Benjamini-Hochberg method (“fdr” method parameter provided to the p.adjust function in R) to obtain the Q-values. Taxa were assigned to different significance tags based on their significance of their associations. The taxa with Q-values <= 0.15 was assigned tag of 3; those with Q-value > 0.15 but p-value <= 0.05 were categorized as 2; the rest were assigned tag 1. Finally, the Association values for each taxon for each of the 18 diseases were computed as a product of directionality (-1, 0, 1) and significance tags (3 or 2 or 1) (**Fig 3A**). Consequently, all taxa-disease associations were categorized as one of -3, -2, -1, 0, 1, 2, 3. Negative numbers indicated taxon was more abundant in control samples than in disease samples and vice versa. After the computation for each of the 196 taxa across the 18 diseases considered for the iteration, the iteration-specific unranked health-association scores were computed for each taxa as:

[Total significant Negative association values (i.e. total number of -3s) – Total significant Positive association values (i.e. total number of +3s)] / [Total number of -3s and +3s]

These unranked health-association scores obtained for the 196 species were the rank-scaled from 0 to 1 (using the same formula as described above) to obtain the iteration-specific health-association scores. The health-association scores for each taxon obtained for the 10 iterations were then averaged to obtain the final taxon-specific Health-Association score.

*Identification of consistent disease-positive and disease-negative taxa*: For this purpose, we also analysed all the 28 clinical conditions together in a single analysis using the same approach. We deduced the set of species that were negatively (-3) and positively (+3) associated with disease. Taxa falling into either the disease-positive and disease-negative categories were those that were significantly (association values of -3 and +3) associated with at least 30% of the total diseases and for whom the total disease-positive (association values of +3) or disease-negative (association values of -3) constitute at least 70% of the total significant associations (association values of -3 and +3). Using the above method we identified 45 disease-negative species and 35 disease-positive taxa.

### Computation of Taxa-specific Health-Associated Core Keystone (HACK) index and microbiome-specific mHACK scores

Taxa-specific HACK index were envisioned as a combinatorial index that combines the association scores obtained for a given taxa with respect to the three properties (core-influence, stability-association and health-association) into a single measure. HACK index combines three scores (computed as previously described) as a product of two values:

- *Average Score*: This is the mean of the three scores (Core Influence: CS, Stability- Association: SS, Health-Association: HS). Higher mean values indicate higher overall association with the three properties. However higher mean values may also originate because of one of the scores showing an especially high value where as others don’t. To address, we calculated the second score called ‘Reward Score’
- *Reward Score*: Reward Score simply indicates how close are the values of the three scores with respect to each other. This was obtained by first computing GINI index across the three values. The GINI index is a measure of the extent to which a given set of values deviate from a perfectly equal distribution. The more similar are the values of a set, the closer will be the GINI index to 0. Thus, GINI index is an inverse measure of similarity within a set and varies from 0 to 1. Consequently, in this case, the Reward Score which measured the extent of similarity was thus calculated simply as: 1 – GINI Index of the three scores. Higher Reward Scores for a taxa indicates higher similarity of the CS, SS and HS. To compute this, we used the Gini function of the DescTools package (v0.99.50) in R

The HACK index for a given taxa ‘I’ was thus computed as

HACK-Index^I^ = Average-Score^I^ X Reward-Score^I^

Higher Hack-Index for a given taxa indicates uniformly high association scores for the taxa with respect to the three properties. In a similar manner, microbiome-level mHACK scores were calculated as the average HACK index of all taxa detected above a certain threshold (to avoid false positive detections; here we used 0.0001 as the threshold).

### Validation of the HACK indices and the mHACK scores in 11 additional cohorts

Details of the 10 additional cohorts have been summarized in **Supplementary Table 17** and provided in detail in **Supplementary Text 3**. Core-influencer taxa for each dataset were identified using the 3R approach (**Fig. 1B**). For a pairwise group comparisons of HACK- indices of core-influencer taxa and non-core-influencer-taxa, we used Mann-Whitney tests using the wilcox.test function of the base R package. The details of the datasets and the methodology adopted for investigating the replicability of the associations of HACK indices with stability and different aspects of health in each dataset are described in **Supplementary Text 3**.

### Validation analysis of the HACK indices with response to Mediterranean Diet Interventions and ICT

*Mediterranean Diet Intervention*: Raw shotgun-sequenced metagenomic reads were downloaded from https://www.ebi.ac.uk/ena/data/view/PRJEB33500. The dietary and anthropometric data were requested and obtained from the authors. Correlations between taxa-specific HACK indices and MED-diet compliance scores were obtained in a two-step manner. In the first step, we obtained the correlations between the individual taxa and the MED-diet compliance scores across samples for each time-point-subject-group scenario. In the second step, these taxa-specific correlations were then correlated with the taxa-specific HACK indices. For investigating the association of HACK-indices to the intake of various dietary components, the intake of each component was first normalized by dividing it by the total energy consumption per subject per time-point. The normalized frequencies were then ranked across all 186 samples (62 subjects X 3 time-points). The association of HACK indices to the intake of each different dietary components were then computed using the same approach as that utilized for the MED-diet compliance scores.

For investigating the correlation between different diet-associated alterations (between week4 and baseline) with microbiome alterations of different species groups, we adopted the following approach. First, we ranked both the taxa-abundance and the normalized dietary-intake profiles across the 186 samples. For each subject, we obtained the dietary alteration and microbiome alterations as the difference between the week4 and the baseline ranked dietary intake (for each component) and ranked taxa abundance (for each taxa) profiles, respectively. Separate microbiome-alteration profiles were computed for the top 18 HACK taxa and the remaining non-HACK taxa. Variations in the dietary as well as different (HACK and non- HACK) microbiome alteration patterns between the MED-diet group and the CONTROL-diet groups were computed by first computing the Principal Coordinate Analysis (PCoA) (using Spearman distance measures) of the alteration profiles, followed by PERMANOVA on the distance matrices. PCoA was computed using the dudi.pco and adonis2 function of the ade4 and the vegan package in R, respectively. Correspondence between the dietary and the different microbiome alterations were computed using Procuste analysis (computed using randtest.procuste function of the ade4 package). Correspondence between the abundance changes of the 18 individual HACKs and the MED-diet associated changes were computed by correlating the abundance alterations of each taxa and the MED-diet alteration centroid the dietary PCoA space.

*Meta-analysis of studies relating microbiome with ICT response*: The 5 studies included in the analysis are summarized in Supplementary Table 18. Meta-analysis of each taxa with ICT response was performed using Random Effects Model as previously described^11^, where in the individual study-specific effect sizes were computed as Hedges’ G between responders and non-responders (with more positive estimates indicating higher enrichment in responders). The HACK-based taxa groups were obtained by dividing the taxa into seven groups G1-G7 (where in the first six groups contained equal-sized partitions of 30 taxa each and the last contained 16 species). Correlations between summarized estimates and HACK scores were computed using Spearman Correlations. Directionality of association of taxa were computed as the difference between the number of cohorts where a taxon was enriched in responders and that where the taxon was enriched in non-responders.

### Identifying the functional/metabolic profiles identified with HACK indices and HACK taxa

*Identification of (experimentally validated) metabolic functionalities associated with HACK indices:* We have previously collated a list of experimentally validated metabolic functionalities for different bacterial and archaeal taxa available as part of different studies^7,18^. These encompassed 361 metabolic functionality profiles pertained to the production, consumption and degradation of various compounds and metal-ions from 990 species. 102 of these 990 species overlapped with our list of 196 taxa investigated for computation of HACK- indices. These experimentally-validated species-to-metabolite-production/consumption mapping encompassed a total of 271 compounds/ions from 990 species. The metabolic functionality profiles were processed as binary matrices of species versus metabolic functionality profiles where the values of 1 indicated a presence of a given metabolic functionality in a given species and 0 otherwise. We then adopted a two-step approach to identify the metabolic functionalities associated with HACK indices and HACK taxa. In the first step, to narrow down our search-space of select features associated with HACK indices, we built Random Forest (RF) models to predict the taxa-specific HACK indices using these metabolic functionalities as predictors. For this purpose, based on the RF output, each metabolic functionality was then a score called the fractional cumulated importance. This was calculated as follows. All metabolic functionalities were arranged in descending order based on their feature importance scores. For a given metabolic functionality (at say rank ‘i’), we then calculated the sum of all feature importance scores from the top feature till the i^th^ given functionality. We then divided this value by the total feature importance scores obtained across all metabolic functionalities. This value was referred to as the Fractional Cumulated Importance score. The Fractional Cumulated Importance basically captures how much of the model performance is retained if we select the top i^th^ features as compared to considering all. Here, we selected a list of 97 of the 361 metabolic functionalities that accounted for 95% of the overall model performance. In next step, we further validated each of these 97 metabolic functionalities for their association with HACK indices using logistic regression models (where we used the detection of a functionality as a response and the HACK index of the corresponding taxa as predictor). The estimate and p-value of this association were obtained for each 97 functionalities. Later, we applied random forest model on these 22 features (p-value <= 0.05) to predict the HACK Score. Interestingly, even as small as 22 features resulted in the Pearson correlation of 0.54 between predicted and actual value of the HACK Score.

*Curation of a high-quality genomes databases species-level genomic functional annotation using various schema:* To obtain the genome-level functional profiles of the taxa investigated here, around 65,000 genomes (including Metagenome assembled genomes) encompassing 121 different core gut microbial species were curated from globally renowned open-source and comprehensive databases like EMBL-EBI MGnify and UNINA MAGs Genome Repository. Among these huge number of genomes, 31913 genomes which are of high quality (with greater than 90 percent completeness and less than 5 percent contamination) were taken for building the framework.

*Genome Annotation*: All selected high-quality genomes were annotated against eggNOG Database 5.0.2 using the latest eggNOG-mapper pipeline (version 2.1.11). The pipeline proteomes corresponding to the genomes were predicted using Prodigal (version 2) which is incorporated inside the eggNOG-mapper pipeline. The annotation files have been leveraged to build the functional profile matrices of the genomes by counting the frequency of occurrences of each functional features in a genome using an R script. The quantitative presence of functional signatures has been captured through these matrices. Profiles of functional features have been generated for eight different categories, those are CAZy (carbohydrate-active enzymes), COG (cluster of orthologous groups), BiGG (Biosynthetic gene clusters), KEGG Orthologs, KEGG Reactions, KEGG Modules, EC number and Pfams.

*Functional Profiles of Unique Species*: From the prior sections it is obvious that multiple genomes have been taken into account for every unique core gut microbial species. In order to synthesize a profile matrix for 121 core human gut microbial species from the profile matrix of 31913 genomes and to assign a single row for each representative species, the arithmetic mean values were computed for each functional feature across all the genomes present for a particular species, thus condensing the multivariate information into a singular summary row representing the profile of a unique species. Thus, profile matrices for eight different functional categories as mentioned above have been generated covering 121 core species.

*Selecting the most informative functional features associated with HACK-indices:* Given the high dimensionality of the data consisting of 47,621 functional features encompassing the eight functional schema (Pfam: 9,807; BiGG: 10,095; cazy: 118, EC: 2,969; COG: 9,370; KEGG Orthologs: 9,617; KEGG Module: 717; KEGG Reaction: 4,928), the first step was to find the filtered set of important functional features associated with HACK indices. Here, we proceeded with only 121 species due to absence of functional profiles of remaining one. To get the subset of important features from 47,621 overall features we followed the RF approach, similar to that followed for the identification of experimentally-validated metabolic functionalities (described above). We first applied RF on functional features from each of the functional schema keeping the HACK Score as response variable. To get the important features from RF model, we took Fractional Cumulated Importance approach as described above. Features were deemed as important whose ratio with the maximum cumulative value was greater than or equal to 0.95. Combination of the important features from different functional profiles generated a set of 6,782 important features (Pfam: 1,674; BiGG: 373; cazy: 54, EC: 750; COG: 1,386; KEGG Orthologs: 1,330; KEGG Module: 284; KEGG Reaction: 931).

*Identification of functional features associated with HACK taxa*: Our next target was to identify the set of features that were associated with the HACK Species. We deduced such features using two approaches where one was based on frequency count of the features in HACK and Non-HACK species and another was based on the gaussian model of the logistic regression. In first approach, we determined the directionality of the feature association with HACK and Non- HACK by taking the difference of mean frequency count of a functional feature in HACKs with that computed for the Non-HACKs. Then we measured the significance of the observed difference using the Mann-Whitney test on HACK v/s non-HACK feature counts. We selected only those features (582) that were significantly positively associated (Q-value<=0.15 and P- value <=0.05) with HACKs. In the second approach, we used logistic regression with the gaussian model to check if the detection pattern of the features was associated with HACK indices (without partitioning species into HACKs and non-HACKs). We shortlisted a set of 584 features that were significantly (Q-value<=0.05) positively associated with the HACK Score. By taking the union of features from the two approaches we got 912 features that were potentially associated with the HACK Score. These features were used to develop the protein database using the proteome information of 18 HACK species.

