## Supplementary Figures for "Towards a Health-Associated Core Keystone (HACK) index for the human gut microbiome"

Author Affiliations

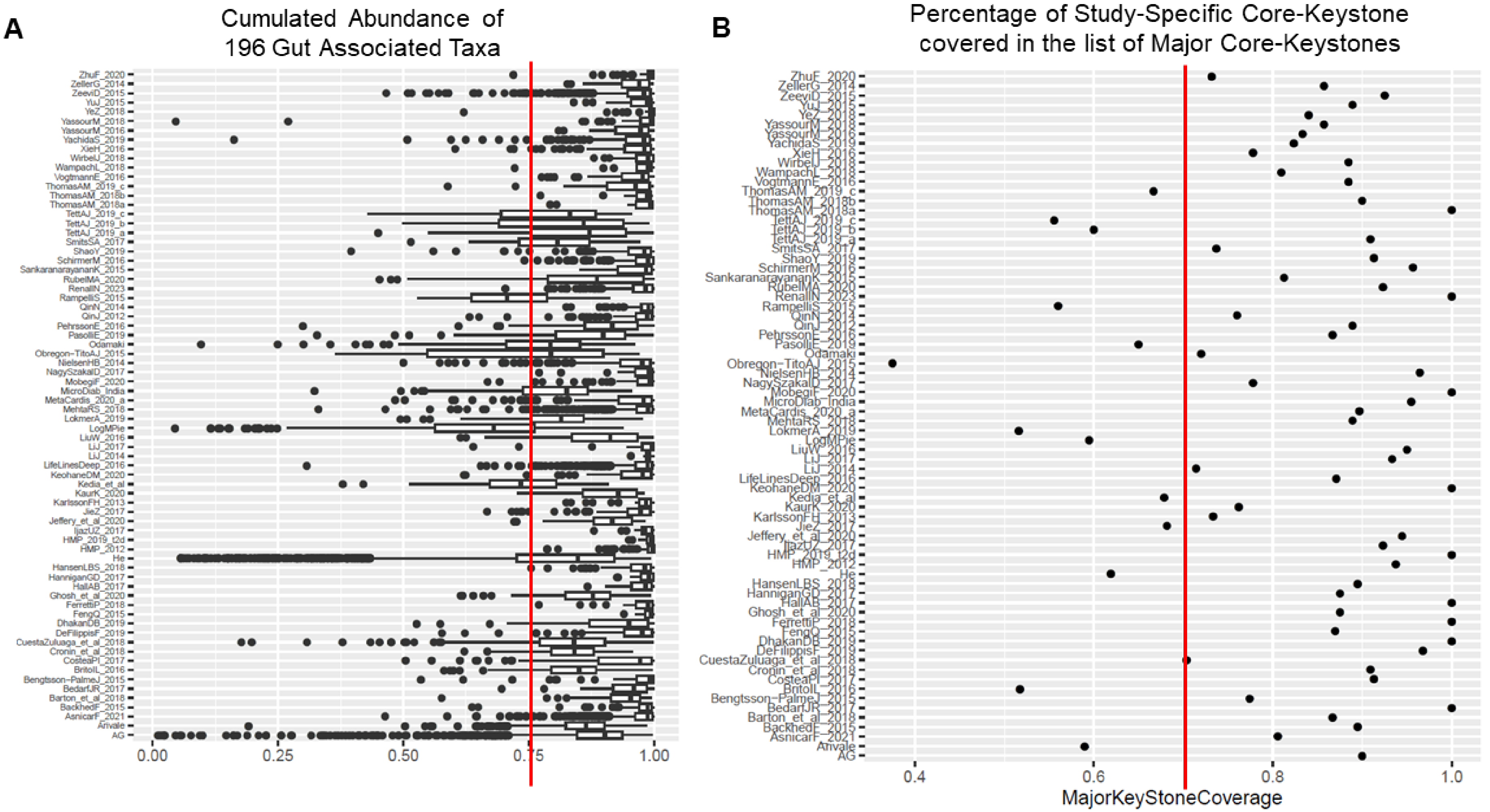


Supplementary Figure 1

**Supplementary Figure 1. A.** Boxplots showing the distribution of cumulative abundances of the 196 gut-associated taxa across the samples comprising each of the 72 study cohorts. These taxa accounted for a median abundance of 97.7% across all 72 studies, with only two studies having a median cumulative abundance < 75%. **B.** Percentage of the study-specific list of Core-Connected-Taxa across the 72 study cohorts that are composed of the top 47 Core-Connected members (as listed in Fig 1C).

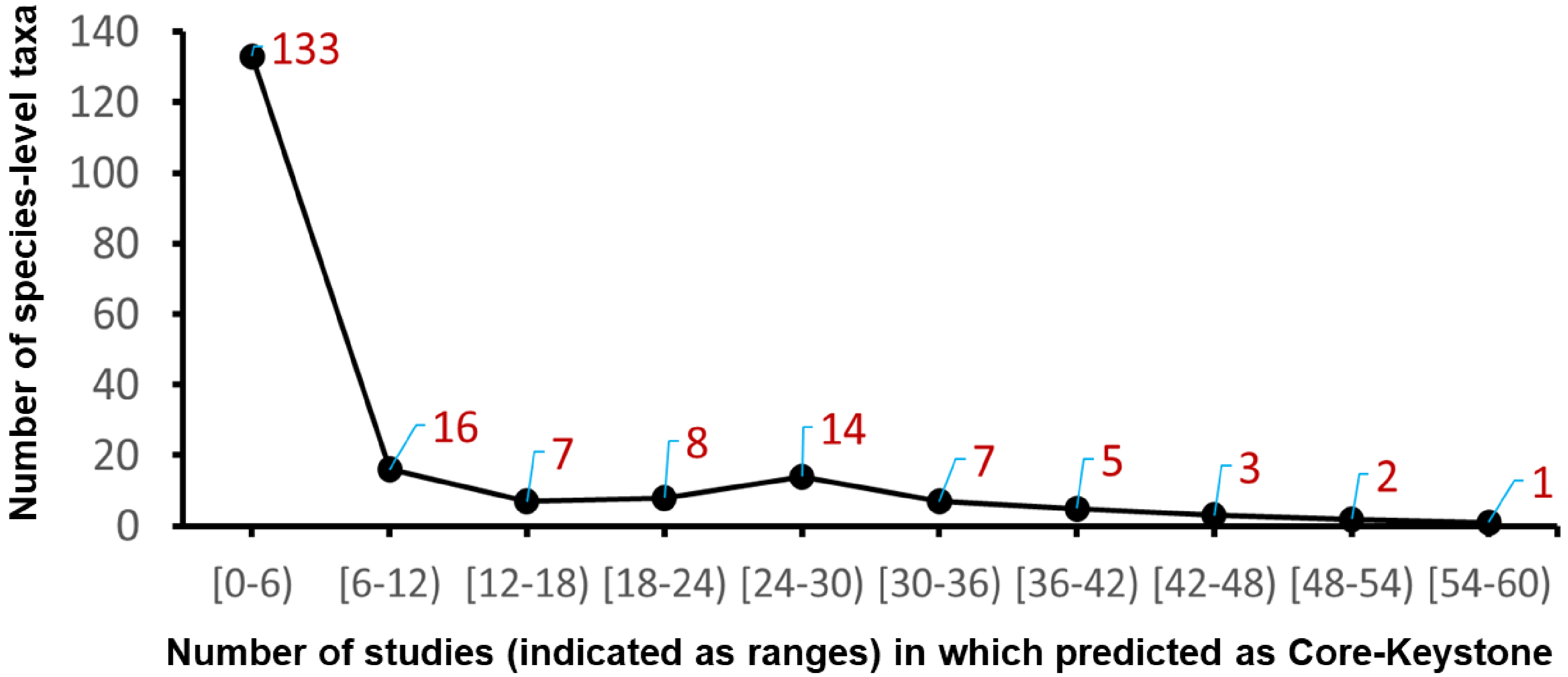


**Supplementary Figure 2**. Detection pattern of the various taxa as Core-Connected members across the various studies. 149 of the 196 taxa were detected as Core-Connected members in less than 12 (of the 72) studies (< 83%). The rest 47 showed a ranked order.

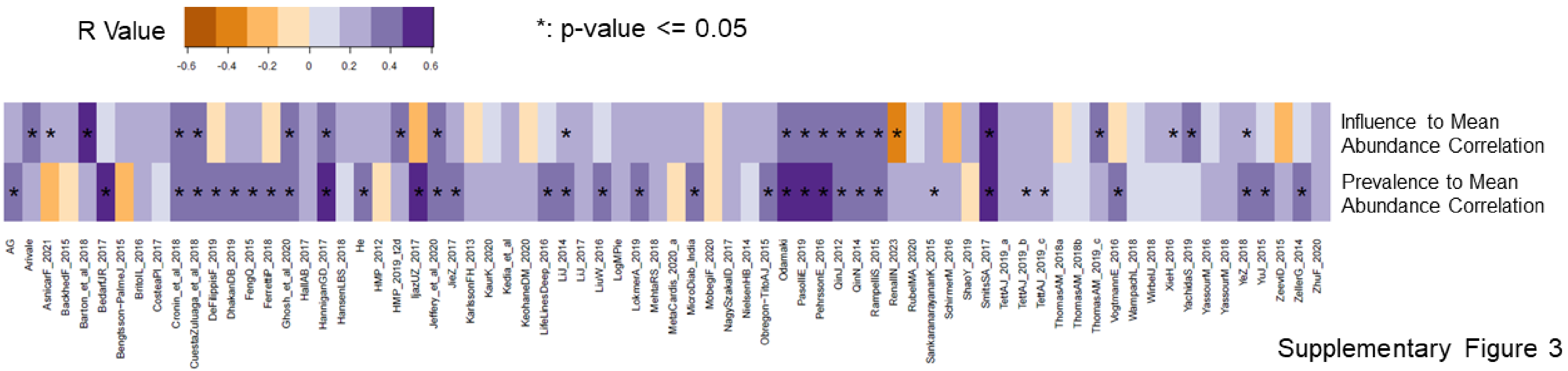


**Supplementary Figure 3**. Heatmap showing the correlation between the Mean Abundance of the 196 taxa with their Influence (computed as the Ranked R-squared) and the Prevalence in the 72 study cohorts of non-diseased adult gut microbiomes. Across all 196 taxa, a significant positive correlation between study-specific mean abundance and study-specific prevalence was observed in only 48% of the cohorts. Similarly, a significant positive correlation between study-specific mean abundance and study-specific influence was observed in only 21% of the cohorts. These observations indicate no clear correspondence between the abundance of a species and its detection as a Core-Influencer member.

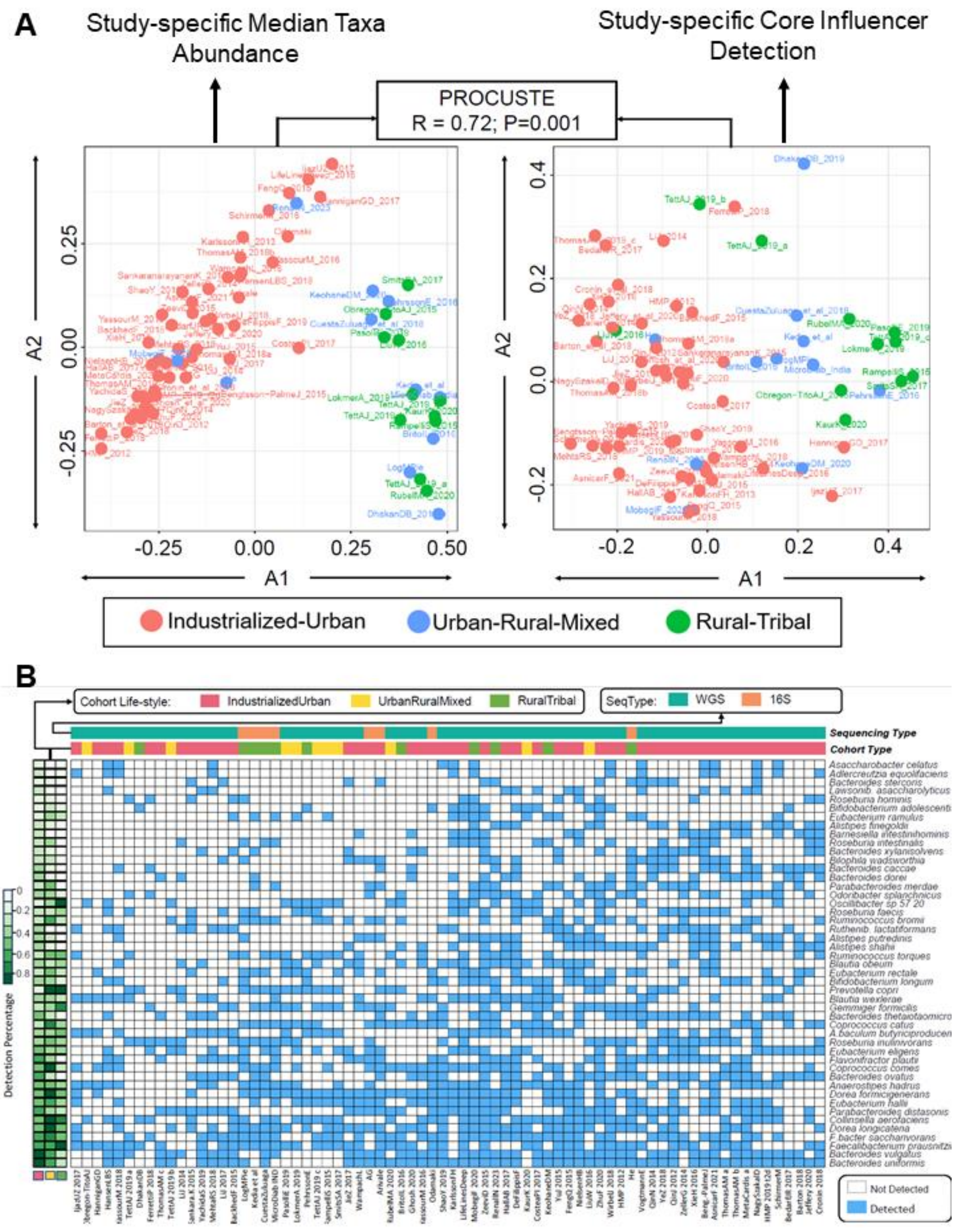


**Supplementary Figure 4. Study-specific variations in the detection of Core-Influencers influenced Cohort-Lifestyle. A.** Principal Coordinate Analysis plots the study-specific variations in the median abundances of the 47 Core influencer taxa (Fig 1C) (on the left) and their detection pattern as Core-Influencers in the different study cohorts. There is a strong concordance between both the study-specific median composition as well as the study-specific detection of the core-influencers (Procuste R: 0.72; P=0.001). This variation is strongly influenced by the life-style of the Study cohorts (classified as either 'Industrialized-Urban',

‘Urban-Rural-Mixed’ and ‘Rural-Tribal’). This variation is further shown in **B**, where the detection pattern of the 47 taxa as core-influencers across each of the 72 study cohorts are shown in as heatmap. The Cohort-Lifestyle as well as the Sequencing Type of the each of the Study cohorts are also indicated. The overall detection percentage of each taxa separately in study cohorts grouped based on Cohort-lifestyle is also indicated as a green heat-strip on the extreme left of the heatmap.

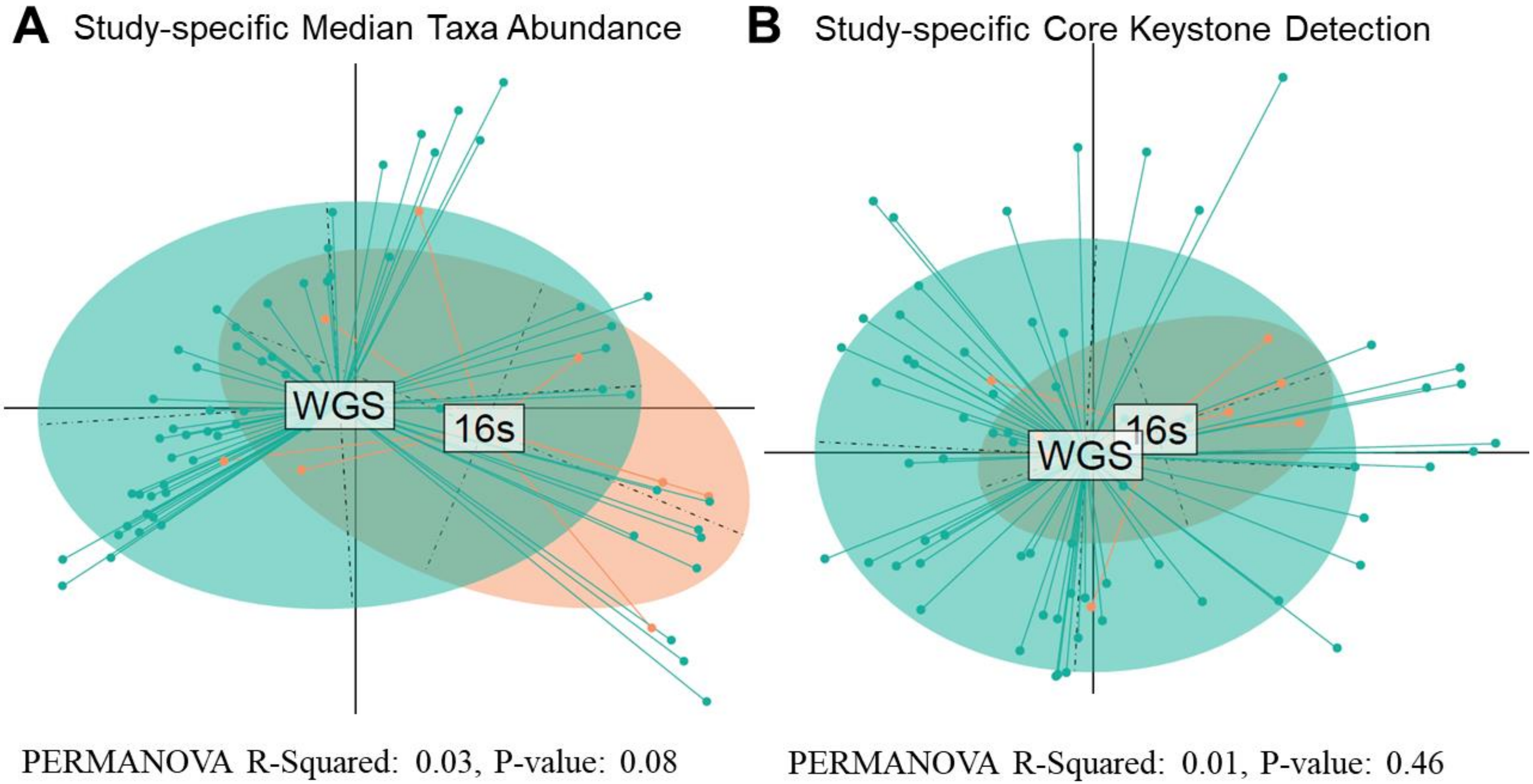


**Supplementary Figure 5**

**Supplementary Figure 5**. Results of the Principal Coordinate Analysis showing the association of **A.** Study-specific Mean Taxa Abundances and **B.** Study-Specific Core-Influencer taxa detection with the Sequencing Type (16S or WGS) of the corresponding Study-cohorts. Also indicated are the PERMANOVA analyses quantifying the associations of the variations of these properties with the sequencing type of the cohorts.

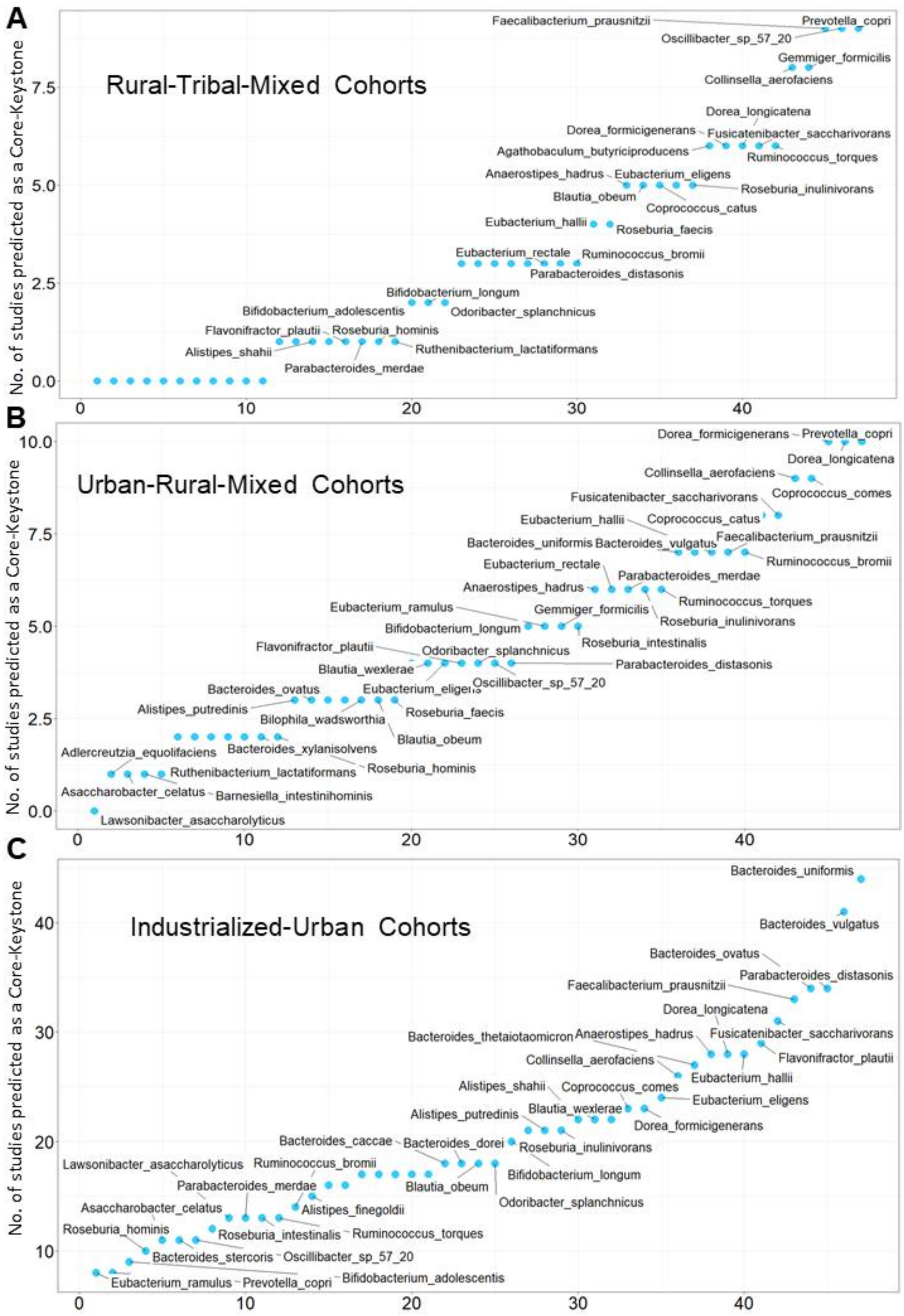


**Supplementary Figure 6**. Ranked order of gut microbial taxa arranged in decreasing order of number of study cohorts they are identified as a Core-Connected members in **A.** Study-cohorts with Cohort-Lifestyle tagged as 'Rural-Tribal', **B.** Study-cohorts with Cohort-Lifestyle tagged as 'Urban-Rural-Mixed', **C.** Study-cohorts with Cohort-Lifestyle tagged as 'Industrialized-Urban'.

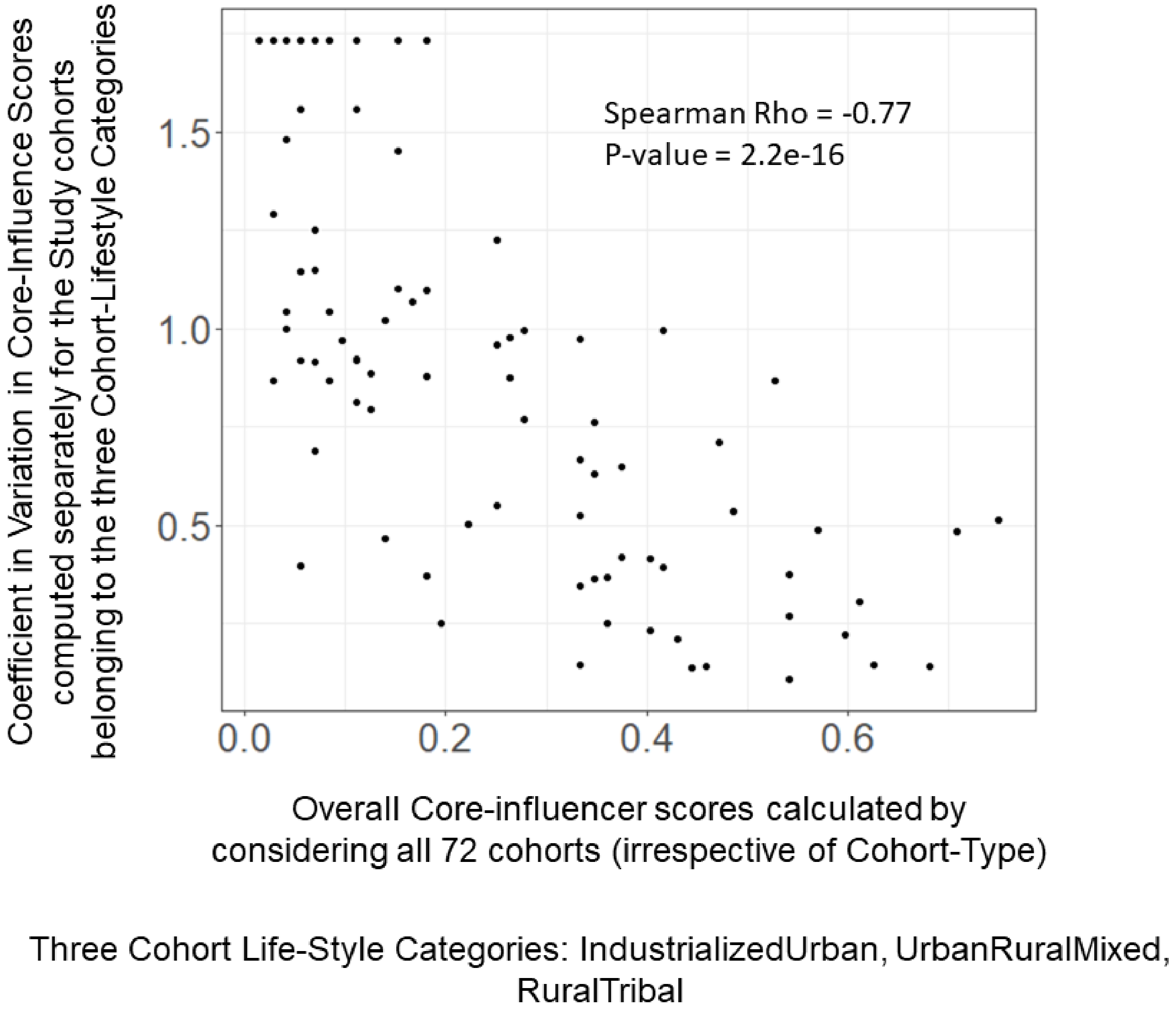


**Supplementary Figure 7**. The more consistently a taxon is detected as a Core-Connected member across the 72 studies, the lower is the variability in its Core-Connectedness across study cohorts from different Cohort-Lifestyle groups. Each point here indicates a species-level taxon. The X-axis shows the percentage of studies the given taxon is detected as a Core-Connected member and Y-axis shows the Coefficient of Variation of the detection rates of the same taxon across the study cohorts belonging to the three different Cohort Life-style categories.

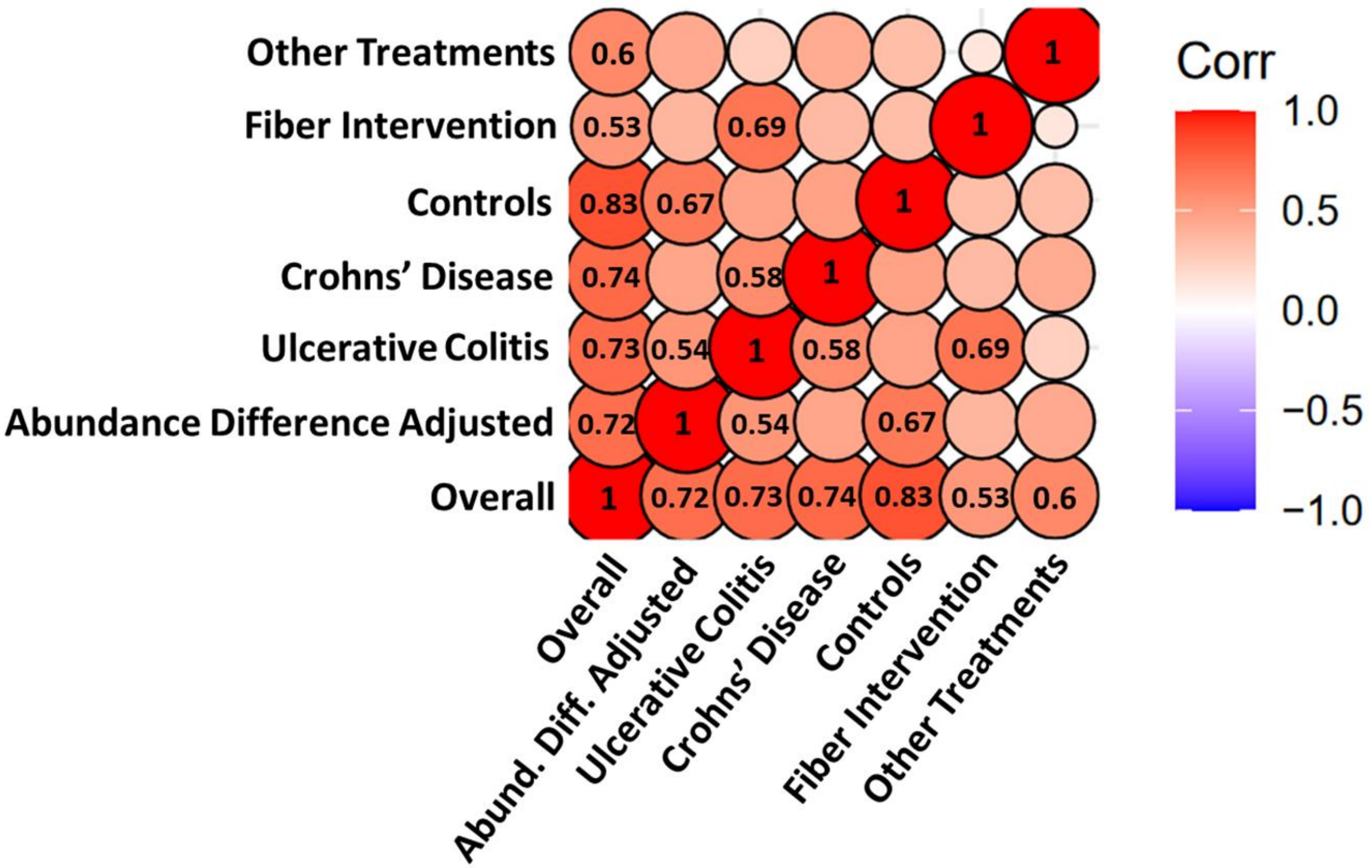


**Supplementary Figure 8.** Heatmap showing the correlations between the species-specific stability-association scores computed using different approaches and subject sub-groups. 'Overall' indicates the SS scores computed overall across all the groups without adjusting for the abundance variations of the corresponding taxa. These were the final scores utilized for the taxa-ranking. 'Abundance Difference Adjusted' indicates the stability-association scores computed for the different taxa, where in the associations of taxa-abundances and follow-up distances were computed after adjusting for the abundance variations of the corresponding taxa across time-points. All other variants (Control, Ulcerative Colitis, Crohns' Disease, Fiber Interventions and Other Treatments) refer to variants of Stability-association scores computed within the different subject groups.

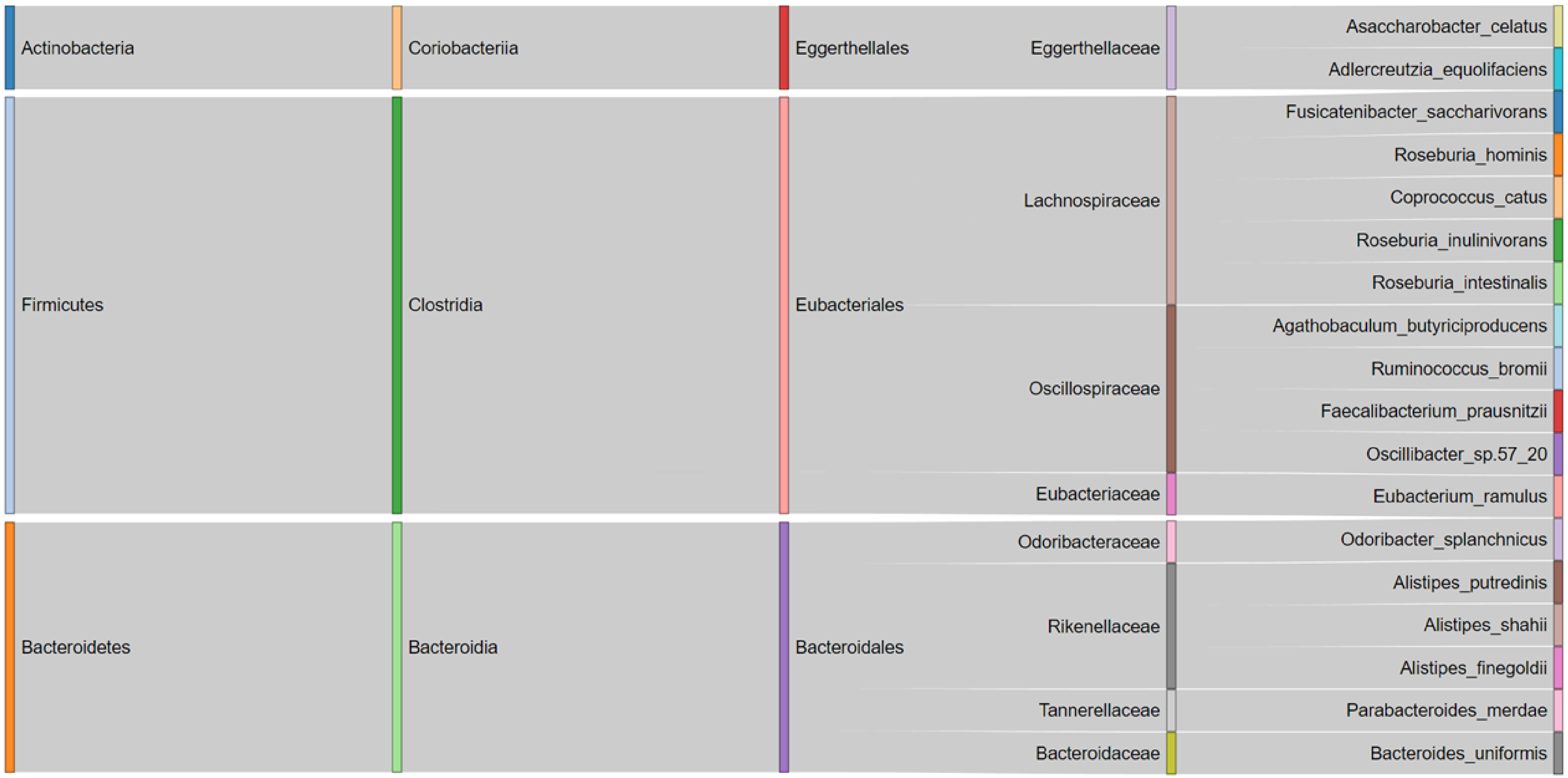


**Supplementary Figure 9.** Complete lineage (from phylum to species level) of the 18 HACK taxa shown as a Sankey plot

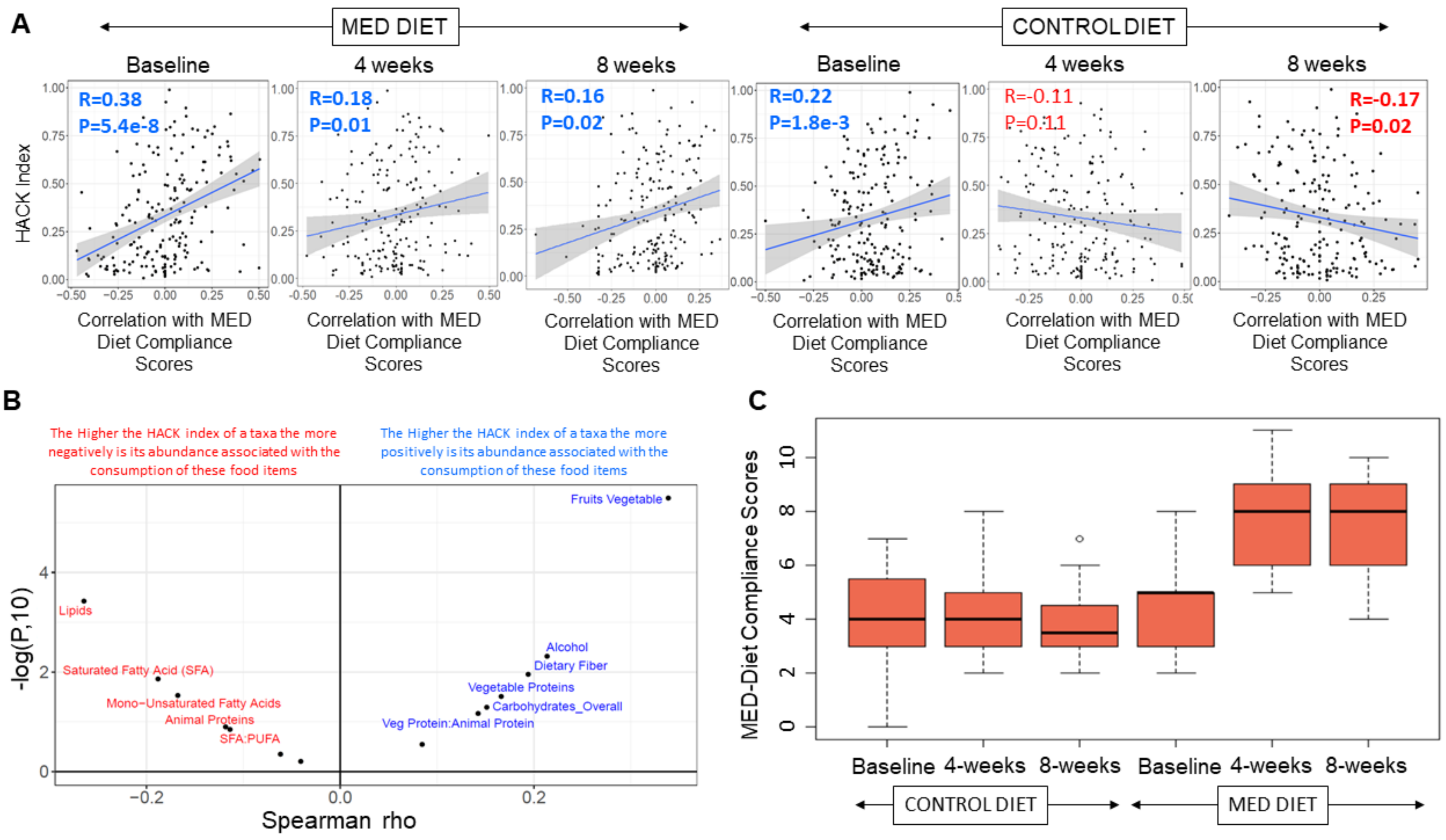


**Supplementary Figure 10**. **A.** Scatterplots showing the correlation between taxa-specific HACK indices and their correlation with MedDiet compliance scores across each of the three sampled time-points and the two subject groups of the Meslier *et al* dataset. For four out of the six time-point-subject-group combinations, we observe significant positive correlation (Spearman Rho and p-values indicated in each scatterplot) between taxa-specific HACK indices and their association MED-diet compliance scores. In other words, the higher HACK index taxa tend to have more positive association with MED-diet adherence. **B.** Shows the associations of the HACK-indices with intake-volume of specific dietary components as volcano plots (Spearman Rho on the x-axis and logarithm-base10 of the Benjamini-Hochberg corrected q-value on the y-axis). It shows that HACK index is positively associated with consumption of Fruits/Vegetables, dietary fiber, vegetable proteins. In other words, higher HACK index taxa tend to show more positive associations with the intake of these food items. In contrast, higher HACK index taxa tend to show more negative correlations with consumption of saturated fat, lipids and animal protein. **C.** Variation of the Italian MedDiet Adherence scores across the three time-points in the two subject groups.

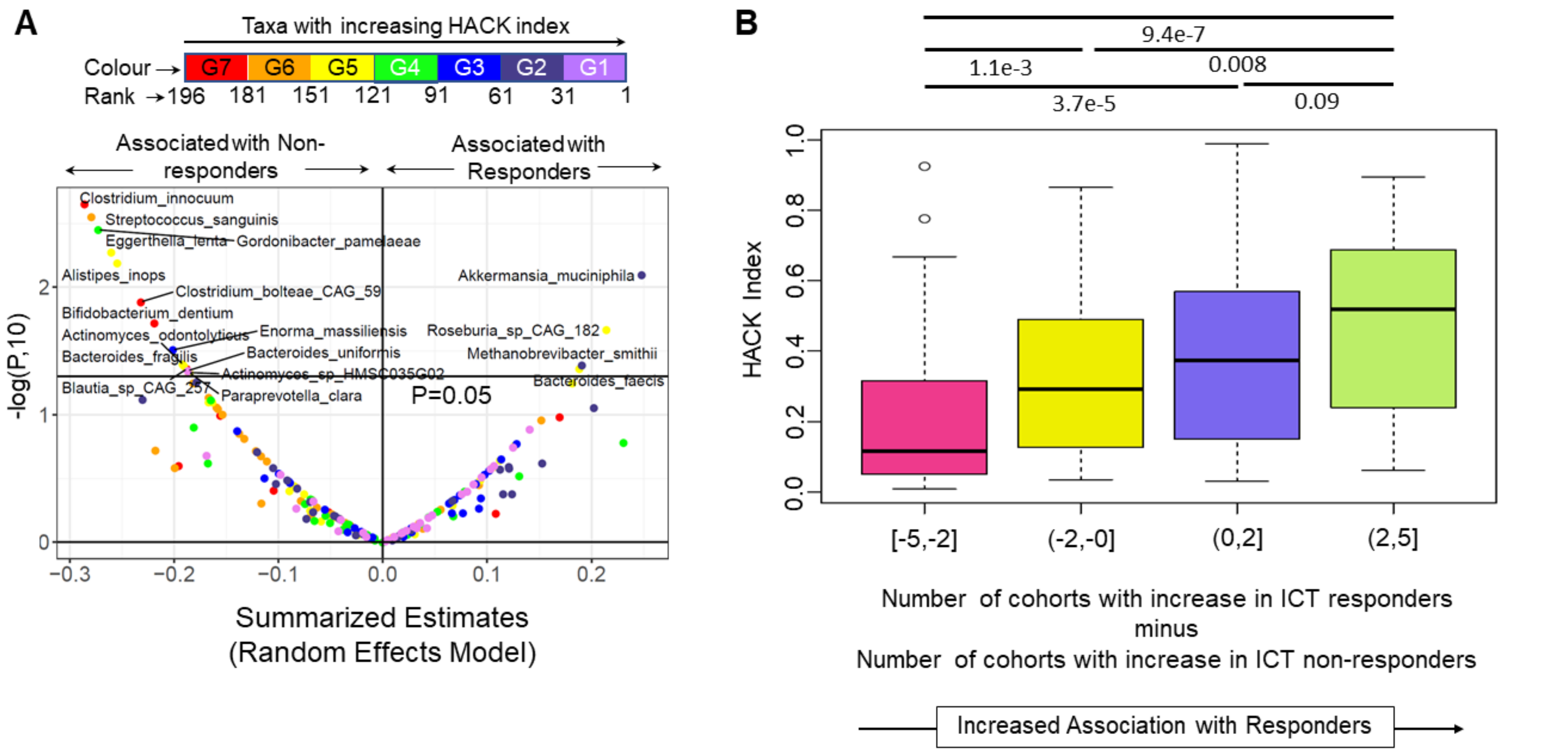


**Supplementary Figure 11**. **A**. Volcano- plot showing the results of the Random Effects Model Meta-Analysis investigating the association of each of the 196 taxa with ICT-response. Taxa showing significant associations with responders as well as non-responders (P<=0.05; Consistency >= 0.80) are highlighted. Taxa-points are coloured based on their HACK-index based groups (or consortia) defined as described in the top panel. **B.** Boxplot showing the significant variations in the HACK indices of taxa grouped based on their consistency of association with responders (calculated as the number of cohorts with increase in ICT responders minus the number of cohorts with increase in ICT non-responders) across the five ICT response cohorts. The higher HACK index taxa show more frequent association with responders.

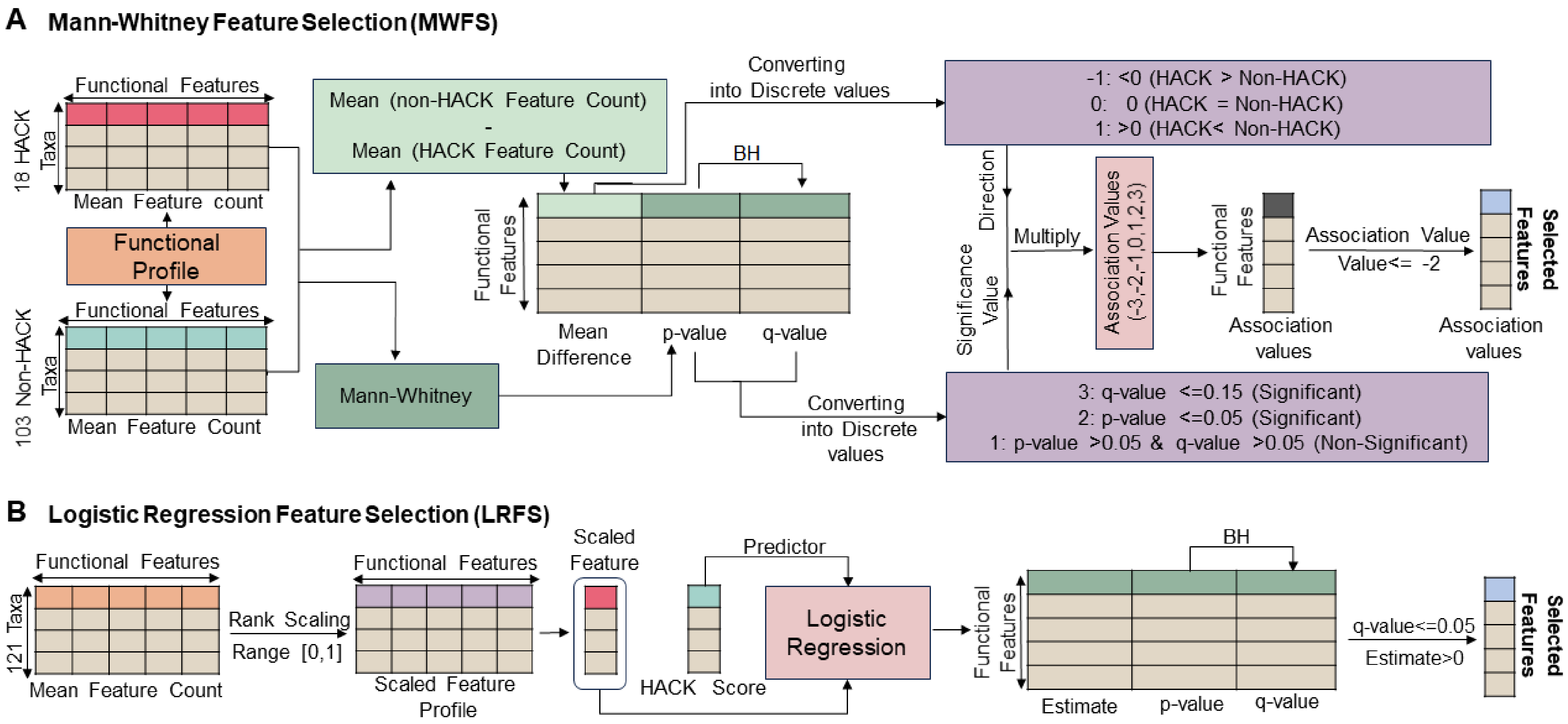


**Supplementary Figure 12**. Pictorial description of **A.** The Mann-Whitney Test based Feature Selection (MWFS) utilized for identifying significantly over-represented in the 18 HACK taxa as compared to the non-HACK ones. **B.** Logistic Regression based Feature Selection utilized for identifying features whose detection rates show a significant increase in taxa with increasing HACK indices.
