## Supplementary Texts for "Towards a Health-Associated Core Keystone (HACK) index for the human gut microbiome"

### Supplementary Text 1

### Details of the collection and processing of the 127 study cohorts utilized for the discovery phase of the study

The distinct subsets of gut microbiomes (and study cohorts) included for the four different investigations are described below:

a. *Stage 1: Computation of Core-Influencer Scores for the gut microbial of 'normal' (apparently non-diseased) adult subjects across a consortium of globally spread study cohorts*: The investigation focused on a total of 18,642 gut microbiomes from apparently 'non-diseased' control subjects collected from 72 study cohorts across 34 countries (**Supplementary Tables 1-2**). Of these, 9,736 (52%) were whole genome sequenced (WGS) microbiome datasets collected from 64 studies. The species-level taxonomic abundance profiles of 58 of these study cohorts (8,922 gut microbiomes) were directly available in the curatedMetagenomicData repository[1]. To this, we added WGS gut microbiome data from six more studies (822 gut microbiomes), corresponding to study populations from Ireland, Australia and the New Zealand[2–6]. To this collection, we further added 8,906 16S-amplicon derived gut microbiome data from eight additional cohorts from US, Japan, China, India and Columbia (including data from the AmericanGut project, the Arivale cohort)[7–13].

b. *Stage 2: Computation of Stability Association Scores using a meta-analytic on diverse longitudinal study cohorts:* We collected a total of 23 longitudinal gut microbiome sequence data as well as corresponding sample metadata from a total of 23 longitudinal study cohorts, conducted in different regions around the world. This combined dataset comprised 9,435 samples obtained from 2,243 individuals. Of these, 7,134 gut microbiome profiles contained at least one follow-up sample from the same subject at a subsequent time-point (**Table 1**). The subjects constituting belonged to different clinical (or phenotypic) categories that were identified in multiple study cohorts, namely 'non-diseased' controls, Crohns' Disease, Ulcerative Colitis, Fiber intervention trials. Additionally, the individuals suffering from difference Colitis sub-variants, (Lymphocytic Colitis, Collagenous colitis), Irritable Bowel Disease, Helminth Infections, individuals harbouring Antibiotic-resistant Enterobacteriaceae, along with antibiotic-treated individuals, fecal microbiome transplantation trials were present in single study cohorts.

c. *Stage 3: Computation of health-association scores using a collection of gut microbiome datasets with matched disease-control samples spanning diverse diseases*: The computation of the health-association scores for the different taxa involved an investigation of 18,020 microbiomes from 62 study cohorts covering 22 nationalities globally (**Supplementary Tables**

**11,12,13**). The selection of this set of gut microbiomes was performed in a step-wise manner as described below.

Initially, we considered 35,447 microbiomes from 114 study cohorts. This contained a total of 14,136 from diseased subjects. These subjects (or individuals) from whom the samples were collected either affected by single or multiple clinical conditions. The metadata for the same were either obtained from the curatedMetagenomicData repository (for those datasets that were derived from this repository) or from the corresponding research papers. We observed that were many related disease conditions that could be further grouped that could be clubbed into disease categories. There were 69 such diseases that were clubbed into 37 disease categories (**Supplementary Tables 12-13**). For each of these disease categories, we computed the study wise control and disease sample counts. Disease-Study combinations which did not contain a minimum of either 15 disease and control-matched samples were discarded. To further ensure that our results are not biased by high variations in the representation of diseased and matched-control subjects, we took only those study cohort for a given disease category for which neither the disease or control-matched samples comprised less than 20% of the total samples. This exercise yielded a final set of 18,020 gut microbiome profiles from 62 study cohorts, encompassing 28 disease categories.

### Supplementary Text 2

#### Association of detected Core-Influencers with the study-specific Sequencing-Type and Cohort-Life-Style patterns

We next investigated if the Core Influence scores of taxa were confounded by any study-specific factors that impact the gut microbiome composition across study cohorts. We investigated two key factors, namely the sequencing method (16S or WGS) and the study-specific geography/life-style patterns (specifically urbanization). We observed that the sequencing type did not have any significant effect on the detection pattern of the core influencer taxa across study cohort (PERMANOVA R-Squared: 0.01, P-value: 0.46) (**Supplementary Figure 4**). To investigate the effect of life-style/geography, we divided the studies into three distinct categories (CohortType), namely 'IndustrializedUrban', 'UrbanRuralMixed' and 'RuralTribal', based on the predominant life-style patterns of the study populations (**Methods**; **Supplementary Table 1**). 'CohortType' was strongly associated with both the microbiome composition profiles and the major Core-Keystone prediction patterns across the study cohorts (**Supplementary Figure 5**) (PERMANOVA: P-value = 0.001 for both profiles). We observed a clear partitioning between the 'IndustrializedUrban' and the 'RuralTribal' study cohorts, with the 'UrbanRuralMixed' interspersed between the two groups. There was a significant concordance between the composition of the gut microbiome and the predicted Core influencer members across the 72 study cohorts (Procuste Analysis: R=0.72; P=0.001) (**Supplementary Figure 5**), and the variation was primarily driven by cohort life-styles.

Besides the previously reported associations of Bacteroides and Prevotella with Industrialized and Non-industrialized populations respectively, we also observed previously unreported cohort-type specific preferences for other taxa (**Supplementary Figure 6**). *Slackia isoflavoniconvertens*, *Oscillibacter sp.*, *Roseburia faecis*, *Gemmiger formicilis*, *Agathobaculum butyriproducens* showed an increasing preference for 'RuralTribal' cohorts and a progressive reduction in Mixed and IndustrializedUrban cohorts, others like *Odoribacter splanchnicus*, *Barnesiella intestinihominis*, multiple *Alistipes* species, *Flavonifractor plautii* and *Bifidobacterium longum* showed preference for the 'IndustrializedUrban' cohorts. Interestingly, we also observed members, like multiple *Coprococcus* and *Dorea* species, *Eubacterium rectale*, *Collinsella aerofaciens* and *Bifidobacterium adolescentis* that showed noticeably high preference of the 'UrbanRuralMixed' cohorts. These distinctive preferences are indications that life-style associated variations in microbiome configurations are associated with preferentially increase in the cross-community associations of certain members.

To further evaluate the extent of impact these cohort life-style specific variations could have on the overall Core Influence scores, we recomputed the Core Influence scores separately for study cohorts belonging to the three CohortTypes. Notably, across the 196 taxa, the coefficient of variation of Core Influence scores (i.e. ratio of the standard deviation divided by the mean of the scores computed individually for the three groups) showed a significant negative correlation with overall Core Influence scores calculated by considering all 72 cohorts (irrespective of CohortType) (Spearman Rho = -0.77, P=2.2e-16) (**Supplementary Figure 7**), indicating that the consistently identified Core-Influencer-Taxa are more likely to be universal.

### Supplementary Text 3

**Investigating the confounding effect of across-time point abundance variations of the investigated and the clinical phenotype of subjects on stability-association scores**.

We further investigated whether the identified associations with stability are not simply driven For this, we performed a second round of Random Effect Model analysis where the cohort-specific effect-sizes were computed between a taxon abundance and the follow-up distances after adjusting for the abundance variation of the investigated taxon between the time-points. The pattern of associations remained unchanged with more than 55% of the significant associations reproduced especially with respect to the Aitchison Follow-up Distances (**Supplementary Table 6**; **Supplementary Figures 8**). In this sub-investigation, while a majority of Core-Keystones retained their significant negative association with follow-up distances (or positive association with stability), the taxa showing associations with an unstable microbiome (positive association with Follow-up distances) included several known 'pathobiont' taxa like *Clostridium innocuum*, *Clostridium hathewayi*, *Clostridium bolteae*, *Ruminococcus gnavus*, multiple Klebsiella spp. along with several oral microbiome associated taxa.

We also sought to determine whether the associations above might be dependent on or confounded variability among the study subjects (mentioned previously; **See Methods**; **Supplementary Tables 7-11; Supplementary Figures 8**). Repeating the above investigation steps individually within the different subjects revealed significant correlations of the stability-association scores computed across each of the subject groups and abundance-adjusted variants with the overall Stability-Association scores computed in **Fig 2A** (ranging from 0.83 to 0.53) (**Supplementary Figure 8**). This suggests that the Stability-Association score robustly captured the overall rank-order of taxon associations with microbiome stability individually within the different subject groups.

### Supplementary Text 4

#### Descriptions of Validation Datasets analyzed for investigating the replication of the association of the HACK-indices with core-influence, stability and health

We investigated a total of 10 gut microbiome datasets corresponding to previously published studies (listed in **Supplementary Table 18**) for validating the HACK-indices computed based on the 127 discovery cohorts summarized in Table 1. The descriptions of the datasets and the various investigations performed on them are summarized below:

**1. *Olsson_16S and Olsson_WGS***: Both the datasets were obtained from a previously published study cohort from Sweden[1]. The datasets were derived from gut microbiome samples collected from a cohort of normal individuals at different time-points throughout a year. The focus of the study was to study the longitudinal dynamics of microbiome amongst a normal group of individuals. From the samples, Olsson_16S and Olsson_WGS were the sequence dataset generated using the amplicon sequencing and shotgun sequencing techniques, respectively. Both datasets were utilized for our investigation of the replication of the stability-association of the HACK-indices. For both datasets (as described in Fig. 2A), we first identified samples with follow-ups (samples collected from the same subject at a subsequent time-point). Follow-up-distances between sample at $T_x$ and $T_{x+1}$ using the Bray-Curtis measure. Across all these samples, the abundances of each detected species with a computed HACK-index were then correlated with the corresponding Bray-Curtis-follow-up distances using Spearman correlation. This was performed using the corr.test function of the psych package v2.3.12 of R. For both datasets, taxa identified to have a significant negative correlation (Spearman Rho < 0 and Q-value <= 0.15) were then listed as the stability-associated markers for each of the corresponding dataset. We then compared the HACK-indices of the stability-associated markers with the non-markers for each dataset.

2. ***Pang***: This dataset was derived from a previously published study from China by Pang *et al*[2]. The study investigated a large cohort from China, totalling around 1575 subjects, including younger individuals to Centenarians. For 45 centenarians, they had also one longitudinal follow-up sample. For this sub-group of 45, we re-performed the stability-association investigation as described above for Olsson_16S and Olsson_WGS datasets.

For this dataset, we also performed an investigation of the abundances of different taxa with declining health in elderly. For the elderly subset (age >= 65 years), the authors had also collected data from the subjects using Life-space Assessment (LSA) questionnaire to stratify patients as either Healthy (H) (LSA >= 66), Less Healthy (LH) (0 < LSA < 66) and Frail (F) (LSA = 0). For our investigation, these categorical groups (H, LH and F) were first converted into ordered numeric variables as H=0, LH=1 and F=2. Subsequently, across the elderly subjects, we then associated the abundance of the different taxa with LSA status (0, 1 and 2) using Robust Linear Regression (RLM) models using the function rlm of the MASS package version 7.3.54 in R. The significance of the associations thus obtained were evaluated using the two-sided robust F-test (using the f.robftest function of sfsmisc package v 1.1.12 in R). Taxa having a negative association (estimate) in the RLM with significance (p-value <= 0.05) were identified to be negatively associated with declining health (or positively associated with health). The HACK-indices of the taxa that were positively associated with elderly health were then compared with those that did not show any associations using Mann-Whitney tests.

**3.** ***Xu***: This was another dataset derived from a previously published study from China investigated by Xu *et al*[3], also focusing on the microbiome and metabolome changes that happen with age in the nonagenarians and centenarians. We included this cohort in our validation analysis to check the replication of our findings of the association of HACK-indices with elderly health observed in the Pang cohort. For the elderly sub-group, the study had measured both physical and cognitive health status using Barthel Score and Mini Mental State Examination, respectively. All individuals with Barthel Score >= 100 and MMSE >= 27 are generally classified to be healthy. We used the same criteria here to identify the healthy elderly (H) individuals. All other individuals in the same group were categorized as less healthy (LH). As for Pang, we converted the categorical H and LH tags as (H=0 and LH=1). The abundance of the different taxa with elderly health status (0 and 1) were then evaluated using RLMs (as described previously). Taxa having a negative association (estimate) in the RLM with significance (p-value <= 0.05) were identified to be negatively associated with declining health (or positively associated with health). The HACK-indices of the taxa that were positively associated with elderly health were then compared with those that did not show any associations using Mann-Whitney tests.

***4. MicroDiab***: This was a dataset derived from a cohort from Denmark previously investigated by Pinna *et al* and Alvarez-Silva *et al* [4,5]. The cohort which was part of a much larger investigation comprising two distinct subject-groups from India and Denmark (the Indian sub-group was included in the discovery cohort), consisted of controls, pre-diabetes (preT2D) and type II diabetes (T2D) patients. As for the above cohorts, we converted the categorical groups of control, preT2D and T2D into ordered numerical variables (Control=0, preT2D=1 and T2D=2). The abundance of the different taxa with disease status (0, 1 and 2) were then evaluated using RLMs (as described previously). Taxa having a negative association (estimate) in the RLM with significance (p-value <= 0.05) were identified to be negatively associated with declining health (or positively associated with health). The HACK-indices of the taxa that were positively associated with elderly health were then compared with those that did not show any associations using Mann-Whitney tests.

***5. Flemer***: This dataset was a derived from a Irish cohort previously investigated by Flemer *et al* [6]. The dataset contained both oral as well as stool samples from Colorectal Cancer (CRC) patients and controls. We focused only on the stool samples. As for the above cohorts, we converted the categorical groups of control, CRC into ordered numerical variables (Control=0, CRC=1). Taxa having a negative association (estimate) in the RLM with significance (p-value <= 0.05) were identified to be negatively associated with declining health (or positively associated with health). The HACK-indices of the taxa that were positively associated with elderly health were then compared with those that did not show any associations using Mann-Whitney tests.

***6. Nagata***: This dataset was a derived from a Japan-based cohort previously investigated by Nagata *et al*[7]. The dataset contained both oral as well as stool samples from Pancreatic ductal adenocarcinoma (PDAC) patients and controls. We focused only on the stool samples. As for the above cohorts, we converted the categorical groups of control, CRC into ordered numerical variables (Control=0, PDAC=1). Taxa having a negative association (estimate) in the RLM with significance (p-value <= 0.05) were identified to be negatively associated with declining health (or positively associated with health). The HACK-indices of the taxa that were positively associated with elderly health were then compared with those that did not show any associations using Mann-Whitney tests.

***7. Song***: This dataset was a derived from a South Korea-based cohort previously investigated by Song *et al*[8]. The dataset contained both oral as well as stool samples from Pancreatic ductal adenocarcinoma (PDAC) patients and controls. We focused only on the stool samples. As for the above cohorts, we converted the categorical groups of control, CRC into ordered numerical variables (Control=0, PDAC=1). Taxa having a negative association (estimate) in the RLM with significance (p-value <= 0.05) were identified to be negatively associated with declining health (or positively associated with health). The HACK-indices of the taxa that were positively associated with elderly health were then compared with those that did not show any associations using Mann-Whitney tests.

***8. Parbie***: This dataset was a derived from a Ghana-based cohort previously investigated by Parbie *et al*[9]. The dataset contained stool samples from Human Immunodeficiency Virus (HIV) patients and controls. We focused only on the stool samples. As for the above cohorts, we converted the categorical groups of control, HIV into ordered numerical variables (Control=0, HIV=1). Taxa having a negative association (estimate) in the RLM with significance (p-value <= 0.05) were identified to be negatively associated with declining health (or positively associated with health). The HACK-indices of the taxa that were positively associated with elderly health were then compared with those that did not show any associations using Mann-Whitney tests.

***9. Saleem***: This dataset was a derived from a Pakistan-based cohort previously investigated by Saleem *et al*[10]. The dataset contained stool samples from T2D patients and controls. We focused only on the stool samples. As for the above cohorts, we converted the categorical groups of control, HIV into ordered numerical variables (Control=0, T2D=1). Taxa having a negative association (estimate) in the RLM with significance (p-value <= 0.05) were identified to be negatively associated with declining health (or positively associated with health). The HACK-indices of the taxa that were positively associated with elderly health were then compared with those that did not show any associations using Mann-Whitney tests.

***Stratifying taxa based on their consistency of associations***: For core-influence, the taxa were classified into three groups based on consistency of identification are core-influencers across the 10 datasets. These groups were those detected as core-influencers <= 10% of datasets (low consistency), 10-50% (medium consistency) and >50% datasets (high consistency). For stability-association, the high, medium and low consistency taxa-groups were identified as those associated with stability in >=2 datasets, single dataset and none of the datasets, respectively. For health-association, we performed nine comparative investigations. The high consistency groups of taxa were identified as those identified to be negatively associated with disease/declining health status in >= 50% (> 4 of the 9) datasets. The medium consistency taxa group comprised those that were detected to be negatively associated with disease/declining health in 10-50% (2 to 4 of the 9) datasets. The low consistency group consisted of all other taxa.

***Comparative investigation of HACK-indices across different consistency groups***: For comparison of mHACK scores across multiple groups, we used the dunn.test implemented in the dunn.test function of the same package (v1.3.5) in R.

### Supplementary Text 5

#### Investigation of the links between HACK-index based ordering of taxa and Mediterranean Diet interventions and Immuno-Checkpoint-Inhibitor therapies

Here, we investigated whether the HACK index-based taxa ordering also correlated with the taxon-specific responses for microbiome-directed therapeutic (dietary) interventions as well as the effects achieved by different taxa-groups in mediating the responses to specific therapies like the immune-checkpoint-inhibitor therapy in cancer.

We first analyzed data from a Mediterranean Diet (MED-Diet) intervention trial previously investigated by Meslier *et al*[1]. MED-diet is a beneficial dietary regime, associated with reduced incidence of several diseases, inflammation and age-associated health decline. Characterized by an increased intake of vegetables, fruits, legumes (and vegetable proteins) and fish and a reduced intake of saturated fats, red meat and milk products, recent studies have shown that the beneficial effects of this diet are driven by an intermediate therapeutic response of the gut microbiome[2,3].

The cohort included 62 individuals, 30 with MED-diet intervention and 32 with habitual-diet, with gut microbiome and diet data profiled at three time-points (baseline, 4-weeks and 8-weeks post-intervention). Across 4 of the 6 time-point-subject-group combinations (exception of 4-week and 8-week time-points in the control-diet group), we observed significant positive relationships between taxa-specific HACK indices and the MED-diet compliance scores of the corresponding subjects (**Supplementary Figure 11A**). Thus, taxa with increasing HACK-indices tended to be more strongly enriched by adherence to the MED-diet. We further validated this MED-diet-linked pattern by investigating the cross-sectional association pattern of taxa with individual dietary components. HACK indices showed a significant positive link with the consumption of dietary components like fruits/vegetables, dietary fibers and vegetable proteins (i.e. the higher the HACK index of a taxa the more positively was its abundance associated with the consumption of these food items), that are positively associated with MED-diet consumption and; a negative link with consumption of lipids, saturated fats and animal proteins (components typically reduced during MED-diet) (**Supplementary Figure 11B**).

Next, we investigated the longitudinal microbiome alterations associated with the MED-diet during this trial. We specifically focused on the alterations between Week4 and the baseline time-points, as these were associated with the strongest alteration of MED-diet compliance scores across the two subject-groups (**Supplementary Figure 11C**). As expected, the subject-groups (CONTROL-diet and MED-diet) exhibited significantly different alterations in their diet intake patterns (PERMANOVA R-squared: 0.18; P=0.001) (**middle panel Fig. 5A**). Notably, the longitudinal (Week4-baseline) alterations of the 18 HACK species not only showed significant different patterns between the CONTROL- and the MED-diet groups (PERMANOVA R-squared: 0.05; P=0.016; **top panel Fig. 5A**), but this alteration pattern significantly correlated with MED-diet linked dietary alterations (PROCUSTE: R=0.28; P=0.001). No such pattern was observed for the remaining non-HACK gut microbiome members. Furthermore, 21/30 (70%) of subjects in the MED-diet group had an increase in the cumulative abundance of the HACK-species between the two time-points (non-HACKs' alteration showing an opposite pattern in these subjects: Fishers' exact test: P=0.019) (**Fig. 5B**). No such pattern was observed for the CONTROL-diet group. We investigated this further at the taxon level by overlaying the directionality of abundance alteration of each of the 18 HACK species with the dietary-alterations associated with the two subject-groups (See Methods). While eight of the 18 HACKs had an increased abundance associated with MED-diet intake, only 3 associated with the change in Diet in the Control sub-group (**Fig. 5C**). Thus, the high HACK-index taxa are positively enriched by increased adherence to the MED-diet. These results indicated that the microbiome response to the MED-diet was predominantly-driven by the top HACK taxa (and specifically associated with an increase of a majority of these taxa), thus validating the role of these taxa as drivers of response to a therapeutic dietary intervention.

We next examined the links between the HACK-based ordering of taxa and the gut microbial determinants of host response to immune-checkpoint therapy (ICT) in cancer patients. A previous cross-cohort investigation has noted the association of the baseline gut microbiome (e.g enrichment of *Akkermansia muciniphila*, *Roseburia*, *Bifidobacterium pseudocatenulatum* in responders) with ICT response, but had also observed that response-associated gut microbial markers were highly cohort-specific and demonstrated limited replicability across cohorts[4]. In this investigation, we investigated five such cohorts for the association of gut microbiome with ICT responses (**Supplementary Table 19**). The objective was to test whether, despite the cross-cohort variability in the association of individual taxa, we could still identify taxa groups (or consortia), that showed reasonably consistent patterns across cohorts and if these groups were related to our HACK-index based taxa order. We performed a preliminary test on the cross-cohort consistency of associations of each taxon with positive ICT response by simply calculating the number of cohorts in which the taxon was enriched (significantly as well as non-significantly) in responders minus the number where it was enriched in non-responders (wherein, the higher the value, the more consistent its associations with responders across cohorts). Here, we observed a clear positive relationship between the taxa HACK indices of the taxa and their consistency of association of responders across the cohorts (**Fig 5D**). This positive relationship was further evident when we divided the 196 taxa into seven groups based on their HACK-indices and performed Random Effect Model based meta-analysis probing the association of each with ICT response across the five cohorts (as summarized in **Supplementary Figure 12A**). Furthermore, validating the previously reported positive associations of *A.muciniphila* and *Roseburia* with ICT response, the investigation also revealed a notable positive link with our computed HACK-indices. The summarized estimates of the Random Effect model showed significant positive correlation with the HACK indices (Spearman Rho = 0.39; P-value=1.9e-5) (**Fig 5E**), indicating that higher HACK indices tend to have stronger overall positive associations with ICT responses. Repeating the meta-analysis on the mean ranked abundances of the seven HACK-index-based taxa groups revealed a significant positive association of the G1 (the taxa-group containing the top HACK-indices) with ICT responders across the cohorts (**Supplementary Figure 12B**). Thus, the top taxa in the HACK-index order (as a consortia) are significantly associated with positive responses to ICT. Notably, the association of the groups also reflected a positive-to-negative association with decreasing HACK indices (with the high HACK groups G1-G4 groups showing association with responders and G5-G7 being associated with non-responders). Our next objective was to identify the functional determinants contributing to the HACK-indices and particularly the functional signatures of taxa with higher HACK-indices.

### Supplementary Text 6

#### Identification of species-level genomic functional tags associated with HACK-taxa

Utilizing the repositories of EMBL-MGNify, NCBI-RefSeq and UNINA-Metagenome-Assembled-Genomes[1,2], we collected 31913 high-quality genomes (completeness > 90% and contamination < 5%) from 121 species-level-taxa (overlapping with our 196 taxa). For each genome, we obtained functional profiles corresponding to 8 different schema (encompassing Biosynthetic Gene Clusters or BiGG, Cluster of Orthologous Groups or COG, Carbohydrate Active Enzymes or CAZy, Enzyme Classification Numbers or EC, KEGG Orthologs or KO, KEGG Modules or KMod, KEGG Reation or KReaction and Pfam, resulting in a total of 47,621 functional features) (**Fig. 6D**). These genome-level function detection profiles were then reduced at the species-level. Subsequently, for each schema, a similar RF-based approach (**as in Fig. 6A**) was used to identify most informative functional features associated with species-level HACK-indices, resulting in a total of 6782 functional features (**Fig. 6E**). As for metabolic functionalities, iterative RF models with 2-fold cross validation (trained on 50% of taxa and tested on the remaining 50%) built using only the informative features indicated strong predictability of HACK-indices based on species-level functional profiles, with high correlations (ranging from 0.5 to 0.6) between predicted and actual HACK-indices observed for all functional schema (except Biosynthetic Gene Clusters: BiGG) (**Fig. 6F**). These observations (especially the validation using the two-fold-cross-validation) suggest that species-level functional profiles are predictive of HACK-indices, can potentially identify newer taxa with high HACK indices.

Encouraged by this observation, we thus proceeded to further probe this select list of 6782 functional features using a combination of logistic regression (**as in Fig. 5A**) (abbreviated as LRFS) and Mann-Whitney tests (abbreviated as 'MWFS'), followed by further filtering based on the detection percentage of the different functional features within the 18 HACK taxa and the detection percentage difference of the same between the HACK and the non-HACK taxa (**Fig. 6G; Supplementary Figure 12**). Using these elaborate steps, we identified a select set of 117 functional features (or tags) with a two-fold increase in the percentage detection in the 18 HACK taxa (i.e. the percentage of taxa in this list, where the tag was identified in at least one of their corresponding genomes) with respect to others and were conserved in >=90% of genomes of >=2 (of the 18) HACK taxa (**Supplementary Table 22**).
